# Combining *Shigella* Tn-seq data with Gold-standard *E. coli* Gene Deletion Data Suggests Rare Transitions between Essential and Non-essential Gene Functionality

**DOI:** 10.1101/038869

**Authors:** Nikki E. Freed, Dirk Bumann, Olin K. Silander

**Author notes:** Corresponding author (OKS).

## Abstract

Gene essentiality - whether or not a gene is necessary for cell growth - is a fundamental component of gene function. It is not well established how quickly gene essentiality can change, as few studies have compared empirical measures of essentiality between closely related organisms. Here we present the results of a Tn-seq experiment designed to detect essential protein coding genes in the bacterial pathogen *Shigella flexneri* 2a 2457T on a genome-wide scale. Superficial analysis of this data suggested that 451 protein-coding genes in this *Shigella* strain are critical for robust cellular growth on rich media. Comparison of this set of genes with a gold-standard data set of essential genes in the closely related *Escherichia coli* K12 BW25113 suggested that an excessive number of genes appeared essential in *Shigella* but non-essential in *E. coli*. Importantly, and in converse to this comparison, we found no genes that were essential in *E. coli* and non-essential in *Shigella*, suggesting that many genes were artefactually inferred as essential in *Shigella*. Controlling for such artefacts resulted in a much smaller set of discrepant genes. Among these, we identified three sets of functionally related genes; two of which have previously been implicated as critical for *Shigella* growth, but which are dispensable for *E. coli* growth. The data presented here highlight the small number of protein coding genes for which we have strong evidence that their essentiality status differs between the closely related bacterial taxa *E. coli* and *Shigella.* A set of genes involved in acetate utilization provides a canonical example. These results leave open the possibility of developing strain-specific antibiotic treatments targeting such differentially essential genes, but suggest that such opportunities may be rare in closely related bacteria.

**Author Summary:** Essential genes are those that encode proteins required for growth and survival in a particular environment. We performed experiments using transposons, genetic elements that disrupt gene function, to determine the set of essential genes in the pathogenic bacteria *Shigella flexneri*. We then compared our results to the well-characterized set of essential genes in the closely related, yet non-pathogenic, bacteria *Escherichia coli*. We found only a small number of genes that are important for growth in *Shigella flexneri*, yet not in *Escherichia* coli. We believe these findings are interesting for several reasons; they help us better understand how quickly the functions of proteins change over time; they suggest possible targets for developing strain-specific antibiotic treatments; and they expand our basic understanding of this pathogen’s metabolic processes.

## Introduction

One general functional characteristic of a gene is essentiality - whether that gene is required for cellular viability and growth. In haploid (e.g. bacterial) genomes, this characteristic can be assessed by attempting to delete a specific gene from a genome. When such a deletion is not possible, this gene is frequently termed “essential” [1], implying that the gene is necessary for cell growth and viability. Gene disruption, although less precise, is more commonly used to infer essentiality using a similar criterion. For example, genes that cannot be disrupted by transposon insertion have been inferred as being essential (e.g. [2]).

One important question is how quickly essential functions change over evolutionary time. If orthologous protein coding genes in two bacterial strains differ in their essentiality classification, this suggests that either the biochemical nature of the protein has changed, or that the cellular context in which the protein acts has changed [3]. It has been experimentally established that such transitions can occur [4-6]. Here we examine how frequently proteins go from being essential to non-essential and vice versa in nature.

A recent study quantified changes in the essentiality classifications of protein coding genes between three alpha-proteobacteria: *Caulobacter crescentus, Brevundimonas subvibrioides,* and *Agrobacterium tumefaciens* [3]. The analysis showed that although orthologous cell components are well conserved, the essentiality of such components (e.g. those involved in the cell cycle) had changed considerably, with only 106 orthologous genes being essential in all three organisms, despite their relatively close evolutionary relationship (89%-93% identity in 16S RNA genes).

In this study we combine dense transposon mutagenesis with high-throughput sequencing (Tn-seq [7]) to quantify gene essentiality in *Shigella flexneri* 2a 2457T (hereafter referred to as *Shigella)*. We compare the essentiality classifications of protein coding genes in *Shigella* with a gold-standard assessment of essentiality in the closely related strain *Escherichia coli* K12 BW25113 (hereafter referred to as *E. coli*) [1]. These two stains are 99.5% identical in their 16S RNA genes and share approximately 70% of their genomic content.

This proximity in evolutionary distance, and the use of a gold-standard data set, brings two unique advantages that have not been available in other studies that have used Tn-Seq or similar methods to quantify gene essentiality [3, 7-19]. First, by relying on the null hypothesis that protein coding genes do maintain their essentiality characteristics, we can objectively assess which quantitative features in the *Shigella* Tn-seq data best predict essentiality or non-essentiality of their orthologous counterparts in *E. coli*; such a comparison to a gold-standard has not yet been used to assess the quality and sensitivity of Tn-Seq data [20], although several studies have validated a small number of Tn-seq-inferred growth defects using clean deletion methods (e.g. [10]). Second, the use of very closely related taxa allows us to quantify on a much shorter time scale the fraction of the essential gene complement that has changed, providing a fine scale window into the rate with which orthologues change in their essential functions.

The data presented here suggest that the essential gene complement of *Shigella* and *E. coli* overlap considerably. Indeed, we find no strong evidence that there are any protein-coding genes that are essential in *E. coli* but not *Shigella*. Conversely, we do find a small number of genes that play critical roles for *Shigella* growth, but which have dispensable roles in *E. coli*, or which are absent entirely from *E. coli*. This implies that the functional correspondence, in terms of essentiality, has changed for only a small number of protein-coding genes.

However, our analysis also suggests that some protein-coding genes that we observe as undisrupted by transposon insertions are in fact not essential for cell growth. Instead, they are either essential for transposon insertion to occur successfully, or their disruption (but not clean deletion) is detrimental to cell growth. This result emphasizes that in high throughput transposon mutagenesis studies, false positive inferences of essentiality may be common, and that simply increasing the resolution or precision of a dataset cannot necessarily solve this problem.

Taken together, our data suggests that the essential gene complement is relatively static over short time scales. However, when protein-coding genes do change from being non-essential to being essential, this appears more likely to occur in pathogenic organisms, perhaps because host environments absolve the organism from manufacturing its own nutrients, or because such organisms have smaller population sizes and are prone to the accumulation of deleterious mutations. It would be interesting to see if this pattern is observed when comparing other pathogens to their free-living sister taxa. If antibiotics can be directed against the function or expression of such differentially essential genes, this may allow targeting such treatments toward specific bacterial strains.

## Results and Discussion

### A Transposon Mutagenesis Library Provides Fine-scale Resolution of Gene Essentiality

We generated a transposon insertion library by transforming a *Shigella icsA* mutant [21] with a plasmid containing a mini-Tn10 transposase with decreased hotspot activity ([22]; **Fig. 1A**, inset) inducible by isopropyl-ß-D-thiogalactopyranoside (IPTG) [23, 24]. After overnight growth on Tryptic Soy Broth (TSB) agar plates containing IPTG, we harvested approximately10^6^ colonies carrying transposon insertions. We pooled and then split this library of clones into six replicates. Three replicates were subject to additional growth step inside the cytoplasm of HeLa cells for four hours. The resulting cells were then harvested, the replicates were bar coded and libraries were prepared. We sequenced all six of these pools on a single Illumina HiSeq lane. For all the analyses presented in this study, we have pooled the data from all replicates and from both of these treatments, as we are focusing on *Shigella* genes that are essential across any permissive growth conditions.

**Fig 1.**
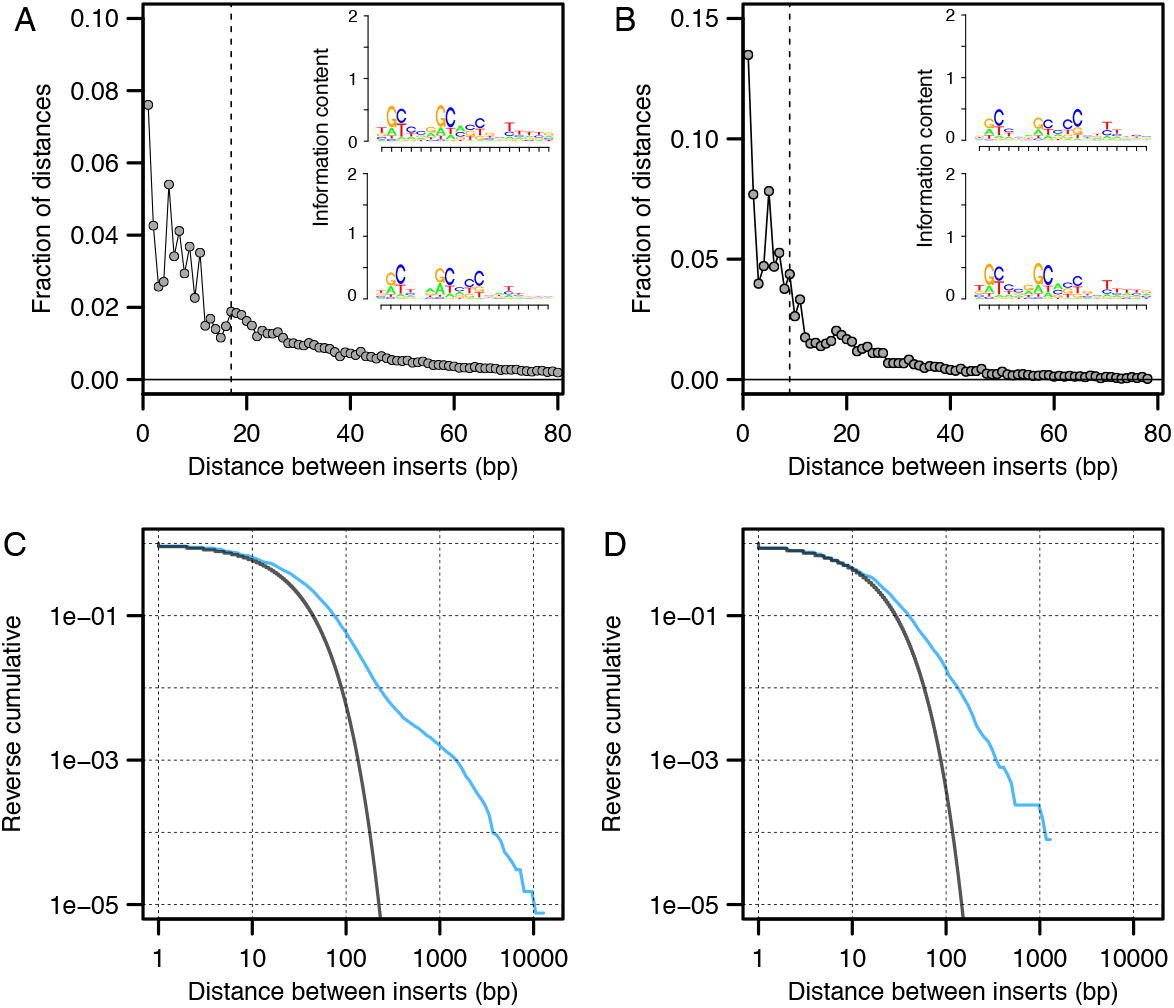
Histograms of distances between inserts on the chromosome for (A) the *Shigella* chromosome and (B) the *Shigella* virulence plasmid. The median distance between inserts is indicated by the dotted line. The insets in (A) and (B) indicate the slight but detectable biases in transposon insert location using a weight matrix motif. The reverse cumulative plots show the observed fractions of distances between inserts for the chromosome (C) and the plasmid (D). In blue, the observed frequencies are plotted. In black, the expected frequencies are plotted, given a geometric distribution (negative binomial with the number of successes set to one) of inter-insert distances (see main text). For both the chromosome and the plasmid, there are considerably more large regions uninterrupted by transposons than one would expect given the geometric null model, observed as a shift of the curve to the right.

From this pool, we mapped insertions at 131,670 unique positions on the *Shigella* chromosome (with many insertions occurring on both the forward and reverse strands but at the same position), and 12,552 unique positions on the large *Shigella* virulence plasmid (see **Methods**). The median distance between inserts on the chromosome was 17 base pairs (bp); on the plasmid this distance was 9 bp. 95% of all inter-insert distances on the chromosome were less than 107 bp; the corresponding figure for the plasmid was 59 bp (**Figs. 1A and B**).

Although the distribution of transposon inserts was relatively even across both the chromosome (**S1 Fig.**), at smaller scales we found many regions in which few or no insertions occurred. Quantitative analyses showed that regions containing no transposon insertions for 100bp or more were considerably enriched (see **Methods**; **Figs. 1C and D**). It is likely that many of these regions are critical for cellular growth in *Shigella*. Indeed, we found that for many of the protein-coding genes in these regions, the orthologous *E. coli* genes are known to be essential (**Figs. 2** and **3**).

**Fig 2.**
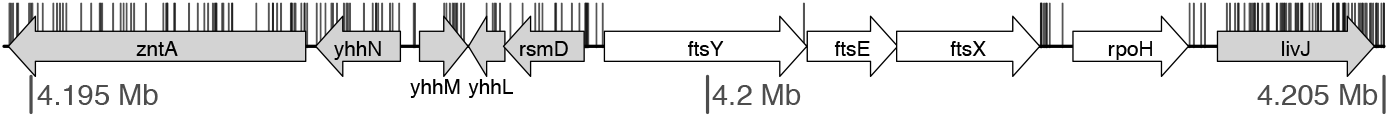
Orthologous genes known to be essential in *E. coli* are also essential in *Shigella*. A region of the *Shigella* chromosome is shown, with genes whose orthologues are known to be essential for growth in *E. coli* (coloured in white) [1, 27], or non-essential (coloured in grey). The unique locations of transposon insertions are plotted as vertical black segments. In the genome region shown here, none of the genes essential in *E. coli* have orthologues that are interrupted in *Shigella*.

**Fig 3.**
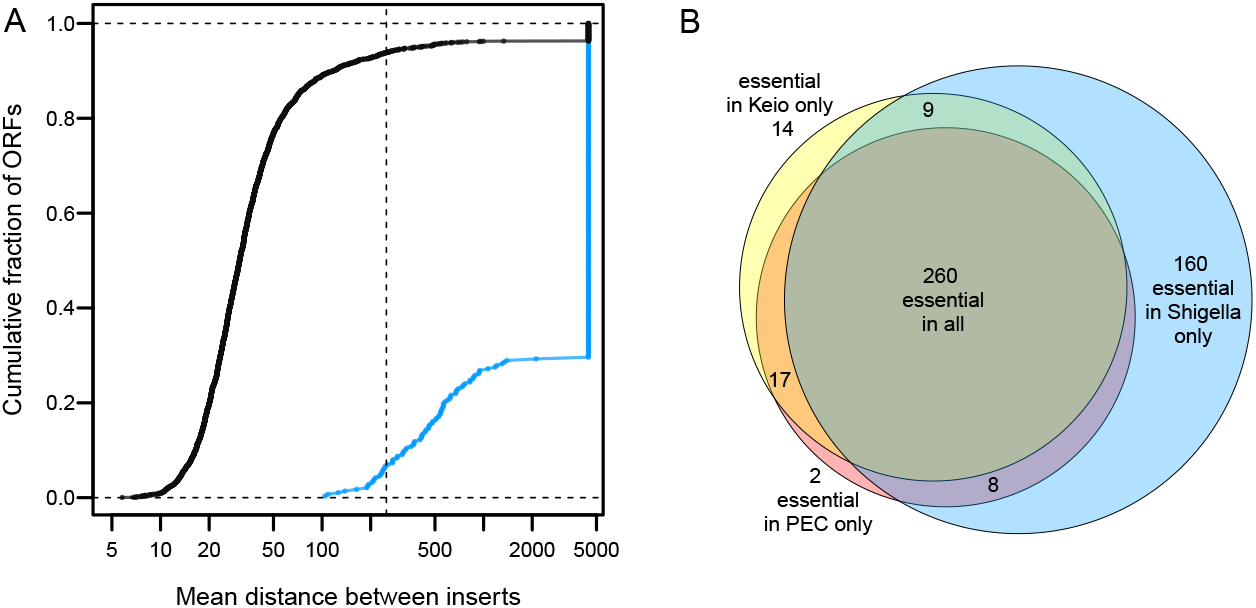
Differences in essentiality classification between *E. coli* K12 and *Shigella*. (A) Cumulative distributions showing the mean distances between inserts for ORFs depending on whether their orthologues are known to be non-essential (black curve) or essential (blue curve) in *E. coli*. All ORFs that are completely uninterrupted by transposons have been plotted at the very right end of the x-axis. The dotted vertical line indicates the cut-off that we used to delineate essentiality in *Shigella* (a mean distance between transposons of 250 bp or more). The 17 blue points to the left of the dotted vertical line indicate ORFs that are essential in *E. coli* but not *Shigella* by our metric. These are likely to be false positives (i.e. non-essential in both *Shigella* and *E. coli)*, as all have inter-insert distances greater than 100 bp (see main text). Black points to the right of the dotted vertical line indicate ORFS that we classify as essential in *Shigella* but not in *E. coli*. Many of these ORFs have *E. coli* orthologues whose deletion genotypes exhibit robust growth, suggesting that their essentiality status has changed. (B) A Venn diagram showing the overlap between essential orthologous ORFs in *E. coli* and *Shigella*.

In contrast to the *Shigella* chromosome, we found that few open reading frames on the virulence plasmid were devoid of insertions. Only six out of 263 plasmid ORFs had no inserts. Two of these were replication proteins (CP0258 and CP0259), and two (CP0217 and CP0218) were located within the plasmid stabilisation region. The remaining two, *mxiH* and *acp*, are both less than 250 bp in length (the cut-off used here to classify ORFs as essential; see below), and thus have a lower likelihood of being hit due simply to their smaller target size. The third replication protein of the plasmid, CP0260, contained a single insert in its 858 bp length. The absence of inserts in the plasmid replication or stabilisation regions is explained by the fact that if such insertions did occur, the plasmid would be lost; such insertions would thus never be sequenced. Thus, this data is consistent with the fact that the *Shigella* plasmid contains no essential genes [25], and suggested that our transposon library provided a fine-scale assessment of which *Shigella* chromosomal ORFs provide critical cellular functions.

### Average Distance Between Inserts Clearly Delineates Essential and Non-essential ORFs

We next quantified which transposon insertion patterns in the chromosome were good predictors of the essentiality of open reading frames. To do so, we first identified 3,027 orthologous open reading frames present in both *E. coli* and *Shigella* for which we also had data on essentiality from both the Keio [1, 26] and the Profiling of the Escherichia coli Chromosome (PEC) [27] studies (**S1 Table**). We considered this combined gene set as a gold-standard of essentiality, for two reasons: it is not subject to artefacts that might exist in Tn-seq dataset, such as insertion biases or biases arising during sequencing library preparation (e.g. [28]); and combining both the Keio and PEC datasets should result in few false positive or false negative essentiality characterizations. This set consisted of 277 orthologues considered essential by both studies, 2,717 genes considered non-essential by both studies, and 33 genes for which the two studies disagree.

We next quantified several characteristics for each protein-coding gene in our Tn-seq data set, including the total number of inserts per ORF, the mean distance between inserts, the length of the 5’ fraction of the ORF upstream of the first insertion, the largest uninterrupted region in the ORF, and others (**S2 Fig.**). We took as a null hypothesis that generally, genes have maintained their essentiality characteristics since the divergence of *E. coli* and *Shigella*. We then tested which of these characteristics best predicted the essentiality status of their orthologous counterparts in the gold-standard dataset of open reading frame essentiality in *E. coli* (the Keio and PEC datasets).

We found that the best predictor of essentiality status in *E. coli* was the mean distance between transposon insertions in their *Shigella* orthologues (Materials and Methods; **S2 Fig.**). For the *Shigella* orthologues of the 277 *E. coli* essential genes, only four had a mean distance between inserts of less than 150bp. 17 (6%) had a mean inter-insert distance less than 250bp. In contrast, only 6% of the orthologues of non-essential *E. coli* genes had a mean distance between inserts of greater than 250bp (**Fig. 3A**). We selected this mean inter-insert distance of 250bp as a cut-off for classifying *Shigella* ORFs as essential, as it provided a balance between protein coding genes classified as essential in *E. coli* but non-essential in *Shigella* (a 6% false negative rate) versus non-essential in *E. coli* and essential in *Shigella* (a 6% false positive rate). By extension, genes that are less than 250bp in length and in which we do not observe insertions were inferred as essential (26 ORFs in total, of which 12 were ribosomal proteins and five were leader peptides). We note, importantly, that almost all of the predictors we tested performed extremely well (**S2 Fig.**).

We next investigated in greater detail the disagreements in essentiality classification between *E. coli* and *Shigella* (**Fig. 3B**). Of the17 *E*. coli-essential genes that this metric identified as non-essential in *Shigella*, all are likely to be false negatives (i.e. in fact essential in *Shigella,* but not classified as such by our criterion). All 17 have a mean distance between inserts of greater than 100 bp (**Fig. 3A**), and nine are uninterrupted for more than 90% of their reading frame. This suggests, surprisingly, that there are no genes that are essential in *E. coli* but whose *Shigella* orthologues are non-essential. This similarity in essentiality is not due to the fact that we use a characteristic that most closely predicts essentiality in the gold standard dataset – this result is robustly corroborated by any meaningful metric that we used (e.g. using other mean distances between inserts as cut-offs for essentiality, using the total number of inserts, the longest uninterrupted gene fraction, or others (**S2 Fig.**)). Overall, this data gives us a very strong prior that genes have maintained their essentiality status (or near-lethal effects on growth) since the divergence of *E. coli* and *Shigella*.

### Many Non-essential *E. coli* orthologues of Essential *Shigella* Genes Exhibit Impaired Growth

As a result of this strong prior, we thus expect that many of the discrepancies in essentiality between *E. coli* and *Shigella* are false positives due simply to the *Shigella* mutants being non-essential, but having significantly impaired growth. Indeed, of the 160 discrepant genes classified as non-essential in *E. coli* but essential in *Shigella*, 34% of the orthologous *E. coli* deletion genotypes exhibit low growth yields (less than 0.5 OD600 after 22 hours of growth in LB [1]). This contrasts strongly with the 2557 ORFs we classified as non-essential in *Shigella:* only 3.7% of the orthologous *E. coli* deletion genotypes had low growth yields (**Fig. 4**). Similar but less striking patterns were observed for growth in glucose minimal MOPS media (**S3 Fig.**).

**Fig 4.**
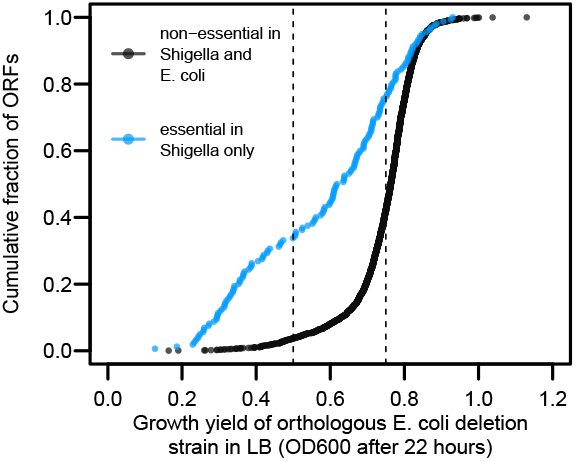
Orthologous gene pairs that are non-essential in *E. coli* but inferred as essential in *Shigella* (blue) tend to exhibit low growth yields in *E. coli*. ORFs that we infer to be uniquely essential in *Shigella* consistently have *E. coli* orthologues with low growth phenotypes in LB media after 22 hours (apparent as a strong leftward shift in the cumulative curve). For genes inferred as uniquely essential in *Shigella*, 34% of the orthologous *E. coli* deletion genotypes exhibit low growth yields (less than 0.5 OD600 after 22 hours of growth in LB). For genes we classified as non-essential in *Shigella* and *E. coli* only 3.7% exhibit low growth yields. Thus, some genes we infer as essential in *Shigella* may not be strictly essential, but instead be required for robust growth. Despite this enrichment for low-growth phenotypes, there are many genes which we infer as essential in *Shigella*, but which have *E. coli* orthologues whose deletion genotypes exhibit robust growth (OD600 greater than 0.75 after 22 hours growth in LB).

It is important to note that Tn-seq assays have only limited power to differentiate essential genes from those whose deletion results in severe growth deficiencies. During the course of preparing the library for sequencing, we estimate that at least 20 generations of growth occurred. If a mutant has a growth rate even 60% that of the wild type, we would expect it to undergo only 12 doublings in contrast to the 20 of the wild type. This would result in a greater than 200-fold underrepresentation of such a mutant (2^12^/2^20^). In addition, this calculation does not take into account any effects that the mutations have on the length of the lag time, which might also have significant effects on the relative frequency of some mutants.

In light of this limited resolution power; given that our prior expectation is that essentiality status changes only rarely; and because we are specifically interested in genes that may have changed in essentiality status, from this point on we focus our analysis on essential *Shigella* genes whose orthologous *E. coli* deletion genotype exhibits robust growth yields (OD600 greater than 0.75 after 22 hours growth in LB (**Fig. 4**)). For these genes, we have relatively high confidence that while their deletion in *E. coli* has few effects on growth, their disruption in *Shigella* is lethal or results in a severe growth deficiency.

### Artefacts of the Transposon Screen Explain Some False Positive Discrepancies

35 essential *Shigella* genes have orthologous *E. coli* deletion genotypes with growth yields higher than 0.75 OD600 after 22 hours growth in LB [1] (**Fig. 4**). Careful inspection suggested that some genes were present in this set due to differences in the growth conditions between *E. coli* Keio and PEC collections and our own. For example, *fhuACD*, and *tonB* all appeared in this set of genes (**Table 1**). All four are involved in iron acquisition, and it is likely that iron was limiting in the solid agar media [29] used during the preparation of the Tn-seq library, as compared to the liquid LB used to measure growth in the Keio study.

**Table 1.** Genes artefactually inferred as essential in *Shigella*.

| Gene(s) | Evidence for Artefactual Inference of *Shigella* Essentiality |
| --- | --- |
| *acrAB*, *tolC*, *ybaB*, *ksgA*, *yebC*, *smpB* (0.73)<sup>1</sup> | Affect kanamycin resistance [30-32] |
| *dnaQ*, *holD*, *recD*, *xseA*, *ruvA* (0.58), *ruvB* (0.60), *ruvC* (0.61), *recB* (0.58), *recC* (0.65) | Likely to affect the transposition process; *dnaQ*, *holD*, *ruvA*, and *ruvB* inferred as essential using Tn-seq in *Salmonella* [13] |
| *priB* | Deficient in plasmid maintenance [33, 34] |
| *fhuACD*, *tonB* | Involved in iron acquisition which is critical for growth in the iron limited media used in this study [29] |
| *rpsT* | S20 ribosomal subunit; new data indicates mutants have poor growth [35] (in conflict with Keio data) |
| *miaA* | tRNA dimethyl transferase; previous data indicates *E. coli* mutants have poor growth [36] (in conflict with Keio data) |
| *ompA* | Outer membrane porin; clean knockouts appear viable [37]; mutant forms are frequently lethal [38] |
| *pitA* | Metal phosphate transporter with ten transmembrane segments; transposon disruption of substrate transporters is three-fold more likely to be inferred as essential compared to clean deletion (see main text; **Fig** ) |
| *potB* | Type I ABC transporter (Putrescine / spermidine transporter) |
| *cysU* | Type I ABC transporter (Sulfate / thiosulfate transporter) |
| *sapB* | Type I ABC transporter (unknown substrate) |
| *ptsH* | Short 306 bp reading frame |
| *ydhR* | Short 258 bp reading frame |

<sup>1</sup> Gene deletions of orthologous *E. coli* genes with growth levels less than OD600 0.75 have these levels in parentheses

For a second set of genes, the discrepancies are likely due to the differences in methodology between the *E. coli* (precise gene deletions) and *Shigella* (transposon inactivation) studies (**Table 1**). We inferred *acrAB* and *tolC* as essential. These genes act together as an efflux pump, and mutations in these genes result in hypersensitivity to antibiotics [39]. Thus, clones with transposon insertions in these genes are unlikely to survive during library growth under kanamycin selection. A similar explanation likely underlies the fact that we inferred *ybaB, ksgA, yebC*, and *smpB* as essential: these four play role in aminoglycoside resistance [30-32].

We also inferred *priB, dnaQ, holD, xseA*, and *recD* as being essential in *Shigella*, although the *E. coli* deletion genotypes exhibit robust growth. All of these are involved in DNA replication, recombination, and double strand break repair, all of which are essential processes in the completion of the transposition process [40]. The related genes *recBC* and *ruvABC* contained a single insert between the five of them, while the *E. coli* deletion genotypes all exhibit only slightly impaired growth of 0.6 OD600 or more (**Table 1**). Certain *recBC* mutants can have considerable effects on the rate of Tn10 excision [41, 42] and we speculate that this may be one reason why we rarely observed insertions in these loci. It has also been speculated that *ruv* mutants inhibit transposition [43]. We propose that after transposition occurs, in order for the event to be successfully resolved, transcription of these genes is often required, and the transposition itself precludes the formation of a proper transcript.

Thus, the dispensability of these ten genes in *E. coli*, and the similarity in their function, suggests that they all affect successful transposon insertion rather than having critical effects on growth. Notably, *priB, dnaQ, holD, ruvA*, and *ruvB* were also inferred as essential in the closely related bacterium *Salmonella typhimurium* via a high-throughput transposon assay. In the same study *ruvC, recB, recC*, and *ybaB*, were inferred as extremely important for growth while *ksgA* and *yebC* were inferred as significantly impairing growth [13]. Again, the majority of these knockouts in *E. coli* exhibit very robust growth (greater than 0.75 OD600 after 22h growth in LB). Given the roles that these genes are known to play in transposition and antibiotic resistance, this suggests that the inference of essentiality may be due to artefacts of the transposon screen.

For a third set of genes, the literature presents conflicting information on the growth phenotypes, with studies that have individually assessed growth rates suggesting poor growth. These include *rpsT* [35], *miaA* [36], and *ompA* [38] (**Table 1**).

There were also two open reading frames that we inferred as differentially essential as they were completely uninterrupted in our data. However, these two open reading frames, *ydhR* and *ptsH*, are very small and less likely to be disrupted, being 306 bp and 258 bp long, respectively. It is probable, then, that this discrepancy is not driven by different physiological roles that they play in *E. coli* as compared to *Shigella*.

Finally, we tested for other possible artefactual patterns in the data based on gene function. We asked whether there were specific functional categories in which genes were more likely to be inferred as essential using the transposon mutagenesis screen in *Shigella* as opposed to clean deletions in *E. coli*. We found two functional categories of genes that showed clear enrichment: genes involved in substrate transport and / or active transport, which were 3- and 2.1-fold enriched, respectively (**Fig. 5**). We hypothesize that one reason for this enrichment is that truncated versions of these proteins disturb the operation of the *sec* machinery, thereby decreasing or stopping growth. Thus, we propose that the four active transporters we infer as essential in *Shigella* but not *E. coli* (**Table 1**) are artefacts due to the transposition process resulting in truncated proteins.

**Fig 5.**
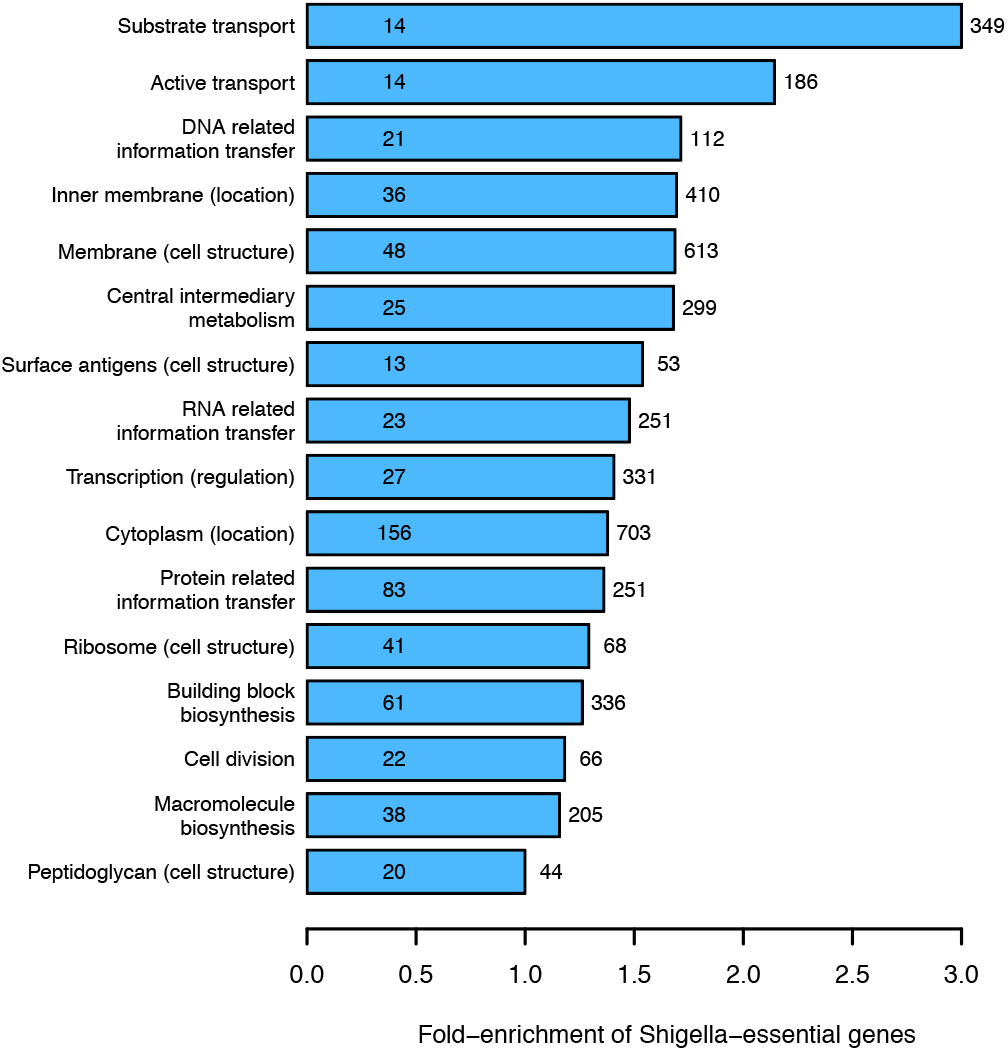
Transposon disruption of *Shigella* genes with transport-related functions are more likely to be inferred as essential compared to clean deletions of similarly functioning genes in *E. coli*. We classified genes according to function using the MultiFun functional classification system [63]. For any category containing more than ten essential *E. coli* genes, we also calculated the number of Shigella-essential genes. As expected, most categories show a relative excess of *Shigella*-essential genes, as we inferred approximately 50% more genes as being essential in *Shigella* versus *E. coli* (**Fig. 3B**). However, two functional categories show a clear excess above this level: substrate transport and active transport, showing a 3- and 2.1-fold increased probability of inferring a gene as being essential in *Shigella* as opposed to *E. coli*. This provides evidence that genes in these functional categories may be more likely to be inferred as artefactually essential. For each functional category (y-axis), we show the number of genes in that category (to the right of each bar); the number of genes found to be essential in *E. coli* (within each bar); and the level of enrichment of essential genes in *Shigella* (x-axis).

### Genes uniquely essential in *Shigella flexneri*

While many differences in essentiality classification between *Shigella* and *E. coli* are likely due to (1) severe growth defects present in both *E. coli* and *Shigella* rather than strict essentiality; and (2) differences in environmental conditions (e.g. iron) between the *E. coli* and *Shigella* assays; and (3) artefacts of the *Shigella* transposon screen that do not occur in the *E. coli* knockout screen, we do find a number of genes which we infer to be uniquely essential to Shigella. We expect that the physiological differences between *E. coli* and *Shigella* are driving these differences in gene essentiality (**Table 2**).

Among the set of genes essential in *Shigella* but dispensable in *E. coli* is *lysS:* this ORF has a functional homologue in *E. coli (lysU* [45]), while in *Shigella flexneri* there is no homologue. Also in this set of genes are *proA, proB, and proC*. These genes act in proline biosynthesis. Given the rich media the cells were grown in, it is surprising that they would be essential. In addition, as *proB* is involved in the first committed step of proline synthesis, its disruption should not cause accumulation of toxic intermediates. However, the data provide strong evidence that the disruption of any these three genes is either lethal or causes severe growth defects (**Fig. 6**). Interestingly, the active proline transporter *putP* is absent from *Shigella* [46]. It is also known that in *Salmonella*, the cryptic proline transporter *proY* is silent [47], and we hypothesize that this may also be true of this transporter in *Shigella*. Thus, inefficient proline transport from the media might necessitate biosynthesis.

**Fig 6.**
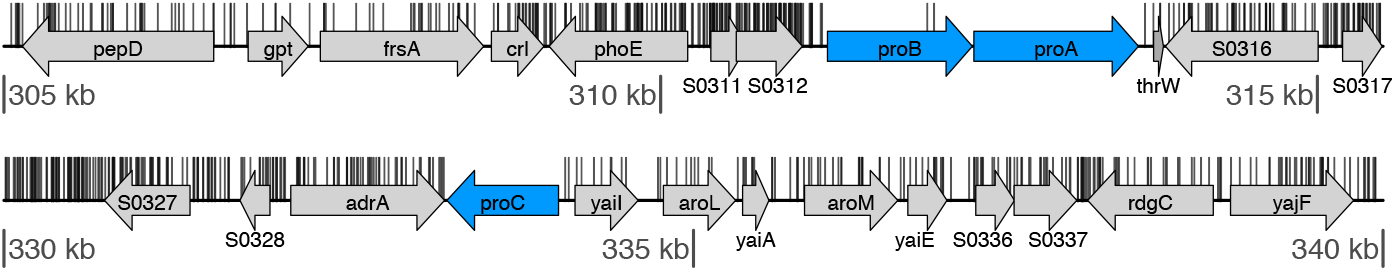
Three genes involved in proline biosynthesis (*proABC*) appear uniquely essential in *Shigella*. The orthologous *E. coli* deletion strains exhibit robust growth (OD600 greater than 0.75 after 22 hours growth in LB), but are essential by our criteria. *proA* and *proC* completely lack transposon insertions, while *proB* contains only two insertions near the 3’end, which leaves approximately 70% of the gene intact, including the entire kinase and substratebinding domain.

A suite of genes involved in acetate utilization (*aceE, aceF, ackA, pta*, and *pykF*) were all inferred as essential in *Shigella* but dispensable in *E. coli*. The significantly detrimental effect on growth that such mutants have has been noted previously using a completely different approach [21]. The difference in essentiality between these two organisms is most likely due to the absence of acetyl CoA synthetase from *Shigella*, and confirms the sensitivity and relevance of our transposon mutagenesis assay for assaying differences between *E. coli* and *Shigella* biology.

**Table 2.** Genes inferred as uniquely essential in *Shigella*. All gene deletions in homologous *E. coli* genes show robust growth in rich media after 22 hours (greater than 0.75 OD600), suggesting that these genes are uniquely essential in *Shigella* as compared to *E. coli*.

| Gene(s) | Function [44] | Evidence for Different Physiological Roles in *E. coli* and *Shigella* |
| --- | --- | --- |
| *lysS* | Aminoacyl tRNA synthetase, tRNA modification | The *lysU* functional homologue is absent in *Shigella* [45] |
| *proABC* | Proline biosynthesis | The active proline transporter *putP* is absent from *Shigella* [46]. The cryptic transporter *proY* may be silent, as observed in *Salmonella* [47], possibly necessitating proline biosynthesis |
| *ackA*<br>*pta*<br>*aceEF*<br>*pykF* | acetate kinase<br>phosphotransacetylase<br>pyruvate dehydrogenase<br>pyruvate kinase | All affect acetate accumulation [48] and utilization [21], which is required for robust growth (*Shigella* lacks the acetyl CoA synthetase present in *E. coli* K12 [49]) |
| *rfbF*, *rfbG*,<br>*rfc*, and *rfbI* | sugar nucleotide biosynthesis for LPS | No *E. coli* K12 orthologues, as this locus has been replaced by the laterally transferred *wbb* locus [50] |
| *spr* | Murein DD-endopeptidase | None known |
| *tufB* | Elongation factor EF-Tu | None known |

For only two other orthologous gene pairs is there strong evidence of discrepant essentiality status: *tufB* (two insert locations; **Fig. 7**) and *spr* (one insert at base pair 543 across 567 bp). For neither of these genes do we have a hypothetical causal explanation. Interestingly, we also found very few transposon insertions in the *tufB* paralogue *tufA* (three insert locations; **Fig. 7**), suggesting that this gene, too, is important for *Shigella* growth despite its relative dispensability in *E. coli* (0.72 OD600 after 22h in LB). We note that these two genes are nearly identical in their sequence, which creates ambiguities in mapping some reads. However, this does not explain the absence of reads mapping to either of them. Understanding the molecular mechanisms driving these apparent disparities in growth phenotypes between *Shigella* and *E. coli* is an important topic for future research.

**Fig 7.**
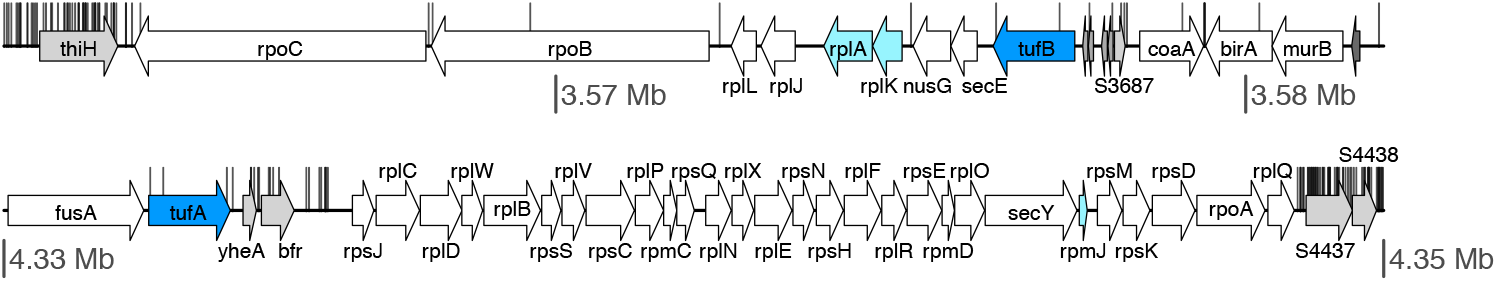
Both elongation factor paralogues *tufA* and *tufB* appear differentially essential in *Shigella* as compared to *E. coli*. The orthologous *E. coli* deletion strains of *tufA* and *tufB* exhibit robust growth (OD600 of 0.72 and 0.78 after 22 hours in LB), but are essential by our criteria. Both genes contain insertions only at the 5’ or 3’ ends of the genes. Genes that are essential in both *E. coli* and *Shigella* are coloured in white. Those inferred as being essential in *Shigella* but for which the orthologous deletion genotypes exhibit robust growth in *E. coli* are indicated in blue. Genes inferred as essential in *Shigella* and which do not exhibit robust growth in *E. coli* are coloured in light blue. tRNA genes are indicated in dark grey.

Finally, the transposon insertion data indicated that a within single large operon, containing the ORFs *rfbACEFGI/ rfc*, four genes completely lacked insertions (*rfbF, rfbG, rfc*, and *rfbl)* (**Fig. 8**). Only *rfbA* and *rfbC* in this operon have *E. coli* orthologues. The remaining genes lie within a commonly laterally transferred region of the *E. coli* chromosome containing *wbbHIJKL, wzxB (rfbX)*, and *glf*. These were all laterally transferred into the K12 lineage [50], replacing the *Shigella-like rfb* operon. The genes in this operon all play a role in sugar nucleotide biosynthesis necessary for O-antigen synthesis and production of the lipopolysaccharide component of the outer membrane [44]. This provides some evidence that specific aspects of this process have become essential in *Shigella*, despite these genes having been replaced by a laterally transferred set in *E. coli* K12.

**Fig 8.**
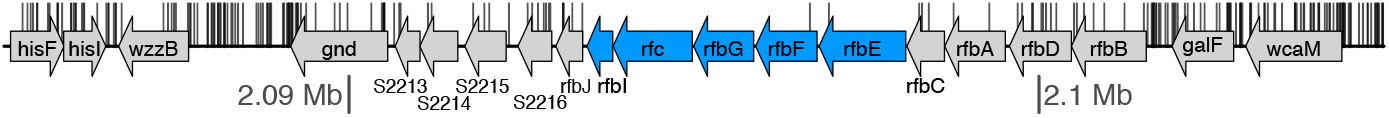
The region of the genome containing the *rfb* operon is largely uninterrupted by transposon insertions. *rfbl, rfc, rfbG*, and *rfbF* are completely uninterrupted by transposon insertions; *rfbE* is uninterrupted over 90% of its length. None of these genes have orthologous counterparts in *E. coli* K12 due to a lateral transfer event that occurred at this locus (see main text). This operon encodes genes active in O-antigen biosynthesis.

## Conclusions

By exploiting the extremely close evolutionary relationship of *Shigella flexneri* with *E. coli* K12, the bacterial strain that has been the most extensively and carefully characterized for its essential gene complement [1, 27], we were able to develop an objective metric to precisely quantify how the results of the Tn-seq data relate to essentiality.

A superficial analysis of our Tn-seq data suggested that a total of 451 ORFs in *Shigella* were essential for cellular growth in rich media. This is very much in line with what other Tn-Seq studies have found, with numbers ranging from 480 in *Caulobacter crescentus* [8] to 447 in *B. subvibrioides* to 372 in *Agrobacterium tumefaciens* [3]. However, it is considerably more than the number that had been found in *E. coli* using in-frame gene knockouts, which is on the order of 300 essential genes. In addition, we found that close to 100% of the reading frames that were classified as essential in *E. coli* K12 were also essential in *S. flexneri*, giving us a strong prior expectation that the essentiality classifications should match between these two taxa.

A more nuanced analysis suggested four explanations for artefactual discrepancies in essentiality between *E. coli* and *Shigella*: (1) many *Shigella* genes were not strictly essential but instead gene disruption caused severe growth impairment; (2) differences in experimental conditions (i.e. iron availability); (3) many of the genes we inferred as essential were important for antibiotic resistance or successful transposition, and are in fact dispensable for growth; and (4) transposon disruption of specific functional classes of genes may result in systematically different effects as compared to gene deletions, for example due to the production of truncated protein products. By carefully dissecting the functions of discrepant genes that do not appear to be artefactual, we were able to pinpoint several genes for which there is some evidence of differential physiological roles in *E. coli* and *Shigella*. Among others, these included *lysS*, three genes involved in proline biosynthesis, and a suite of genes involved in acetate utilization (**Table 2**). In addition to these, we found one large operon which appears to have an essential role in *Shigella* growth but which is missing completely in *E. coli*. Surprisingly, we found only two additional genes that are differentially essential (*tufB* and *spr*) (**Table 2**).

Even after attempting to decrease false positive inferences of gene essentiality in *Shigella*, it appears to be considerably more common for genes to be dispensable for growth in *E. coli,* but critical for growth in *Shigella*. We suggests that one reason *Shigella* more may have a larger complement of essential genes than *E. coli* is that it frequently lives as an intracellular pathogen, and may have lost some of the functional redundancy that is present in *E. coli*. This may occur because host environments provide an abundance of nutrients, or because pathogens requiring a small infectious dose, such as *Shigella* [51], have inherently smaller population sizes and are more subject to genetic drift. A third possibility is that changes in gene function or redundancy may have occurred through selection for increased virulence, which has resulted in the inactivation of certain genes being selectively advantageous. Finally, we note that the discrepancies in essentiality between these two bacteria may be exploited to develop antibiotics that have strain-specific effects [21].

## Methods

### Strains

For all experiments, *Shigella flexneri* 2457T *ΔicsA* was provided by M. B. Goldberg was used as the parental strain. This strain is unable to exploit the host actin cytoskeleton for motility and spreading [52]. Bacterial cells were grown in Tryptic Soy Broth (TSB) media. For experiments using eukaryotic cells, HeLa cells were cultured in DMEM supplemented with 10 mM Hepes, 25 mM glucose, and 4 mM glutamine. *Shigella* were grown to exponential phase in tryptic soy broth, coated with poly-L-lysine, and added at a multiplicity of infection of 25, resulting in an infection rate of around 60%. *Shigella* was centrifuged onto HeLa cells (600 × g for 5 min). At 30 min postinfection, we added gentamicin (100 μg/mL) to kill extracellular bacteria. Bacterial cells were allowed to grow within HeLa cells for a total of 4 hours.

### Transposon library

Using a Tn10 transposon with a T7 promoter [23, 24] we created a library consisting of approximately 10^6^ clones. This library was created by mating *E. coli* strain BW20767 containing the pJA 1 transposon plasmid with a spontaneous nalidixic acid resistant clone of *Shigella flexneri* 2457T Δ icsA for 5 hours. Transposase expression was induced by plating onto TSB plates containing 0.2 mM isopropyl-β-D-thiogalactoside (IPTG). Colonies were allowed to grow at 37°C for 18 hours on TSB agar plates. All colonies from these plates were then pooled and 100μl aliquots of the transposon library were stored at −80°C.

Three replicate experiments were carried out on different days in which an aliquot of the transposon library was grown for 18 hours in TSB to stationary phase, diluted 1:100 and grown to exponential phase (0.7 OD600). This exponential phase culture was split into two: part of the bacterial culture was pelleted and saved and other was used for infecting HeLa cells (as described in [21]). After 3.5 hours, HeLa cells infected with the *Shigella* transposon library were trypsinized and pelleted. Uninfected HeLa cells were also collected and used to spike the original bacterial culture not used for HeLa infection in order to account for HeLa DNA. All resulting DNA was extracted using the Bacterial Genomic Miniprep Kit (Sigma).

### Sequence library construction and sequencing

To amplify the transposon region from these pools, we used one top strand primer annealing to the transposon and a pool of three bottom strand primers each of which consisted of 10 random nucleotides followed by a pentamer of common nucleotides in *E. coli* [53]: N_10_GGTGC, N_10_GATAT, and N_10_AGTAC, using Phusion pfu (**S4 Fig**). A nested PCR was then performed to add the P7 and P5 Illumina adapters, as well as a barcode. The products from this second PCR were then size selected for inserts between 200bp and 300bp, quantified using a Qubit, and sequenced on an Illumina HiSeq2000 at the D-BSSE Quantitative Genomics Facility resulting in 49bp single end reads. We used a custom sequencing primer on the P5 end of the molecule such that on both ends of the molecule, reads started directly on the chromosome.

### Read mapping

In total, we obtained 198,682,954 reads. We found that the number of reads at each location in the genome varied by up to four orders of magnitude. For this reason, we considered only whether an insertion had occurred at a specific location, and not on the number of reads we obtained at a specific location, which is likely to be highly biased due to PCR artefacts. We thus first deduplicated the reads using *tally* [54], and then used bowtie2 [55] to align the reads to the *Shigella flexneri* 2a 2457T genome and the *Shigella flexneri* 2a str. 301 plasmid pCP301. The sequence of the *S. flexneri* 2457T plasmid is not available. However, the *S. flexneri* 2457T and 2a str. 301 plasmids are nearly identical in sequence (differing by 30 SNPs; see below). Sequence reads were not trimmed for quality as read quality is taken into account in bowtie2. We used the --sensitive-local option to allow soft clipping on the 3’ end of the reads (so that reads that contained adapter sequences at the 3’ end could map successfully), and required at least 22bp of matching sequence at the 5’ end of the read.

We checked for single nucleotide polymorphisms (SNPs) on both the chromosome and the plasmid using the samtools mpileup and bcftools utilities [56, 57]. We retained as possible SNPs only those sites that fulfilled the following three criteria: (1) the SNP was inferred as homozygous (necessarily true, as *Shigella* is haploid); (2) the quality score was above 20; and (3) at least three reads on both the reverse and forward strands confirmed the SNP. We found 99 SNPs on the chromosome (as compared to the reference *Shigella flexneri* 2457T in NCBI) and 30 SNPs on the plasmid (as compared to the *Shigella flexneri* 2a str. 301 plasmid in NCBI (in addition to 12 and 2 small indels, respectively). These are detailed in **S2 Table** and **S3 Table,** respectively.

Within chromosomal protein coding regions, 44% of all SNPs were synonymous, while 32% fell outside of genic regions (i.e. protein coding or RNA genes). These fractions are greater than one would expect if such SNPs were randomly located on the genome. Only 24% of all mutations in chromosomal coding regions are expected to be synonymous (not accounting for mutational biases), and only 28% of the chromosome is annotated as nongenic (including repeat regions, although for many of these regions, the absence of an annotation may be erroneous). Additionally, only 2 of the 12 (17%) small chromosomal indels fell in coding regions. This suggests that there was some selection against nonsynonymous substitutions that occurred during the culturing and derivation of the *Shigella flexneri* 2a 2457T *virG* mutant. More importantly, the small number of SNPs that we found suggests that few, if any, reads remained unmapped due to sequence differences between the strain used in our experiments and the sequenced GenBank strain.

In total, the reads mapped to 89,028 unique locations on the forward strand and 83,074 on the reverse strand of the chromosome, for a total of 172,102 insertions. Some of these insertions occurred at identical positions but on opposite strands, so in total, insertions occurred at 131,670 unique sites in the chromosome. Correspondingly, the reads mapped to 8,208 unique locations on the forward strand and 8,585 unique locations on the reverse strand of the plasmid, for a total of 12,552 unique sites. During the insertion of the Tn10 transposon, a 9 bp target DNA sequence is duplicated [58]. We accounted for this duplication in calculating the distances between insertions (by moving the inferred site of insertion for one direction (we arbitrarily selected the antisense direction) backward by 9 bp). Similarly, this duplication was accounted for in calculating various statistics of insertions within genes: sense insertions that were inferred as occurring in the last 9 bp of a gene were ignored in calculating the mean number of insertions per gene (as these bp are duplicated upstream of the insertion). Antisense insertions occurring in the first 9 bp of a gene were ignored, as these bp are duplicated downstream of the insertion.

Using the read frequencies at all unique insert locations, we found that the transposon insertions occurred in a biased manner, integrating more often at sites similar to the known 9bp consensus NGCTNAGC [58], although this bias was relatively weak (**Figs. 1A and B**, insets). This low level of bias is likely due to our using a transposon with reduced hotspot activity [22]. In addition, we found that insertion frequency was slightly influenced by nucleotides further downstream of this 9bp consensus (**Figs. 1A and B**, insets). Sequence logos for this analysis were visualized using the R package seqLogo [59].

The median distance between inserts was 17 bp in the chromosome and 9 bp in the plasmid (**Fig. 1B**), suggesting that the transposon libraries yielded a relatively fine-grained map of the essential genomic complement for both the chromosome and the plasmid.

As expected given the variation in insertion densities across the chromosome, we found high variance in the distribution of inter-insert distances. The total length of the *S. flexneri* genome is 4,599,354 bp in total. Given that we observed 131,670 inserts, under a model of random insertion, we would expect a median distance between inserts of 35bp, with 95% of all inter-insert distances being less than 107 bp (under the assumption that these distances are distributed in a geometric manner (i.e. a negative binomial with the number of successes set to one). For the plasmid, we observed 12,552 inserts over 221,618 bp, such that we expect a median distance of 18bp between inserts, and that 95% of all inter-insert distances are less than 59 bp. However, as noted above, we found that on average transposons insertions were separated by a median of 17bp on the chromosome and 9bp on the plasmid. Fitting a geometric distribution to the observed data over 99% of the range of the inter-insert distances (i.e. from 1 to 237 bp for the chromosome and from 1 to 78 bp for the plasmid) more exactly quantified this over-dispersion, and showed that uninterrupted regions in the chromosome greater than 100 bp were considerably enriched (**Fig. 1C**).

### Paired end read mapping and inference of IS element dynamics

We used 100 bp paired end Illumina sequencing data from this same library to look for structural rearrangements due to IS elements in the genome. However, this analysis was complicated by the fact that many IS elements share close to 100% identity with others around the genome. During these analyses we thus restricted our searches to regions of the genome for which we had *a priori* expectations that they harboured a rearrangement (i.e. if there were no inserts and the orthologous *E. coli* locus was non-essential or absent). Specifically, we followed the following procedure: we extracted a 50 kilobase pair (Kbp) region from the genome surrounding each hypothesized rearrangement (in all cases, this was a deletion). We then used bowtie2 with the paired end option, allowing up to 10 Kbp inserts to map all reads from our 100 bp PE dataset. From these mapped reads, we retained only read pairs that had (1) mapping quality scores greater than 20; (2) at least one read that matched perfectly (i.e. at all 101 bases of the read) to the genome; and (3) were unique in their length at any specific location (thereby excluding artefacts such as PCR doublets). From these paired reads we then inferred the insert size, which is plotted in **S5 Fig**. The vast majority of insert sizes ranged between 100 and 400bp. However, some were much larger (e.g. up to 9,000 bp in **S5B Fig.**). We inferred that these surrounded regions of the genome that must have been deleted.

Such deletions would result in the set of genes contained within as being inferred as essential because of their lack of transposon insertions. However, in the vast majority of cases, we found that when large operons lacked insertions but had non-essential orthologous operons in *E. coli*, or were missing entirely from *E. coli*, these operons were in fact missing from the *Shigella* clone that we used, most likely due to the rapid dynamics of IS elements in this bacterium [60]. For example, no sequence reads we obtained mapped to the *yeaKLMNOP* operon, which spans a total of 9,240 bp. Upon further analysis using a paired end genomic data set, we found that this region was clearly missing from our *Shigella* clone (**S6B Fig**.). This was similarly true for several other operons, as well as for single genes. We did not consider any region in which we identified a deletion in our downstream analyses.

### Essential open reading frames

We identified 3,027 unambiguously ORFs that were present in both *E. coli* and *Shigella flexneri* 2457T [61], and for which we had essentiality data. We used reciprocal shortest distance [62] to find orthologues, with the requirement that the alignment of the two hypothetical orthologues extend over at least 60% of the longer ORF. To establish a gold-standard set of essential genes we combined the data from two studies of the effects of gene deletion on growth in *E. coli* K12: the Keio collection [1] and the PEC study [27]. We retained only those ORFs which we had data on essentiality from both studies. We then quantified which transposon insertion patterns that most closely corresponded with the essentiality delineations in theses studies. Specifically, we selected the feature that maximized the number of true positive essential genes (maximizing the sensitivity) while minimizing the number of FP (maximizing specificity) (this metric is a receiver operator characteristic for which we quantified the area under the curve (AUC; **S2 Fig.**)). We selected from eleven nonindependent features: (1) the total number of insertions; (2) the mean number of bp between insertions; (3) the median number of bp between insertions; (4) the number of bp in the 5’ end preceding the first insertion; (5) the number of bp in the 5’ end preceding the first insertion relative to the total bp in the gene; (6) the number of bp in the 5’ end preceding the second insertion; (7) the number of bp in the 5’ end preceding the second insertion relative to the total bp in the gene; (8) the number of bp in the longest uninterrupted stretch of the gene; (9) the number of bp in the longest uninterrupted stretch of the gene relative to the total length of the gene, and (10) the number of bp in the longest stretch of the gene interrupted by at most one insertion; (11) the number of bp in the longest stretch of the gene interrupted by at most one insertion, relative to the total length of the gene.

We found that for both the PEC dataset and the Keio dataset, the two best predictors of essentiality were the mean distance between inserts (AUC = 0.972 for the PEC dataset, 0.952 for the Keio dataset, and 0.973 for the genes on which both datasets agreed on the essentiality classifications); and the fraction of the gene that lay in the longest uninterrupted region (AUC = 0.969 for the PEC dataset, 0.955 for the Keio dataset, and 0.971 for the genes on which both datasets agreed on the essentiality classifications) (**S2 Fig.**). We selected mean distance as on average, it marginally outperformed the other statistic on the gold standard data set.

We note that for eight of the 14 genes classified as essential solely in the Keio dataset, the orthologous *Shigella* ORFs have mean distances less than 30 bp, suggesting that these genes may be falsely annotated as essential in the Keio study. In contrast, nine of the ten genes inferred as essential solely in the PEC dataset have mean distances greater than 200 bp; the tenth has a mean distance of 189 bp.

### tRNA disruptions

We found insertions in 27 out of 99 tRNAs, with tRNAs for certain amino acids being considerably overrepresented (**S2 Table**).

### Additional analyses of differentially classified essential genes

We also tested for the enrichment of certain functional categories in the set of genes that were classified as being essential in *Shigella* but not *E. coli*. This differs from the analysis present in **Fig. 5** in that we are asking whether across a broad set of functions, are specific categories enriched for Shigella-essential genes. In **Fig. 5** we ask whether *within* a single functional category, is there a much higher fraction of *Shigella* essential genes than we would expect, given the fraction of genes in that functional category that are essential in *E. coli*.

Thus, for this analysis, we separated genes by primary functional category and secondary subcategory using the MultiFun designations (e.g. the primary functional category cell processes divided into the secondary subcategories of cell division, SOS, stress, protection, and motility). We then calculated the fraction of *Shigella*-essential genes within each secondary subcategory and compared this to the total fraction of *Shigella*-essential genes within the primary category (e.g. we calculated the fraction of essential genes in Ribosomal Function (the secondary category) and the fraction of *Shigella*-essential genes in all other categories in Cell Structure (the primary category) (**S6 Fig**.). We tested for enrichment (depletion) using a Fisher exact test.

We also examined gene conservation. Highly conserved genes were considered to be those present in more than 50% of all gamma-proteobacteria [61]. We found that genes classified as uniquely essential in *Shigella* were much more conserved across gamma-proteobacteria (79% highly conserved) compared to genes that were found non-essential in both *E. coli* and *Shigella* (36% highly conserved; p=1.0e-33, Wilcox rank sum test).

### Availability of supporting data

All read data are in the SRA with accession numbers XXX.

## List of abbreviations used

bp: base pairs
*Shigella* – *Shigella flexneri*: 2a 2457T
*E. coli* – *Escherichia coli*: BW25113
ORF: open reading frame
PEC: Profiling the E coli Chromosome database

## Authors’ contributions

NEF, DB, and OKS conceived and designed the transposon mutagenesis. NEF performed the mutagenesis and sequencing. OKS analysed the data with input from DB and NEF. NEF and OKS wrote the paper.

## Acknowledgements

The authors thank Luise Wolf for input on designing the sequencing protocol.

## Endnotes

## Table and Figure legends

**S1 Table**. Full table of gene characteristics and orthologue relationships used in the analyses.

**S2 Table**. List of chromosomal SNPs and indels observed in the *Shigella* strain used here that differ from the GenBank sequence NC004741.

**S3 Table**. List of plasmid SNPs and indels observed in the *Shigella* strain used here that are different from the GenBank sequence of the *Shigella flexneri* 2a strain 301 virulence plasmid pCP301 (NC_004851).

**S4 Table**. Table listing tRNA genes and the number of insertions in each.

**S1 Fig.**
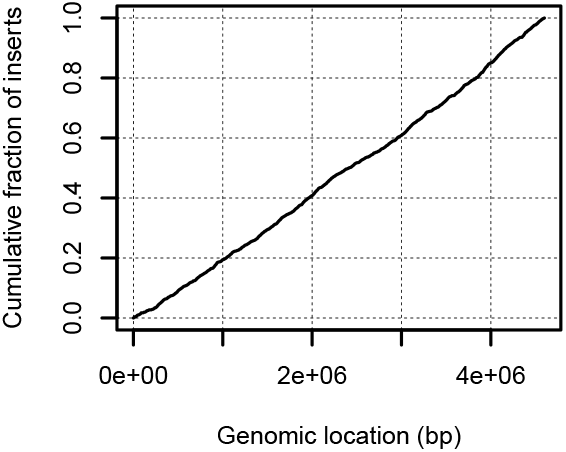
Distribution of transposon insertions across the genome. We observed little bias on the chromosomal level of insert locations.

**S2 Fig.**
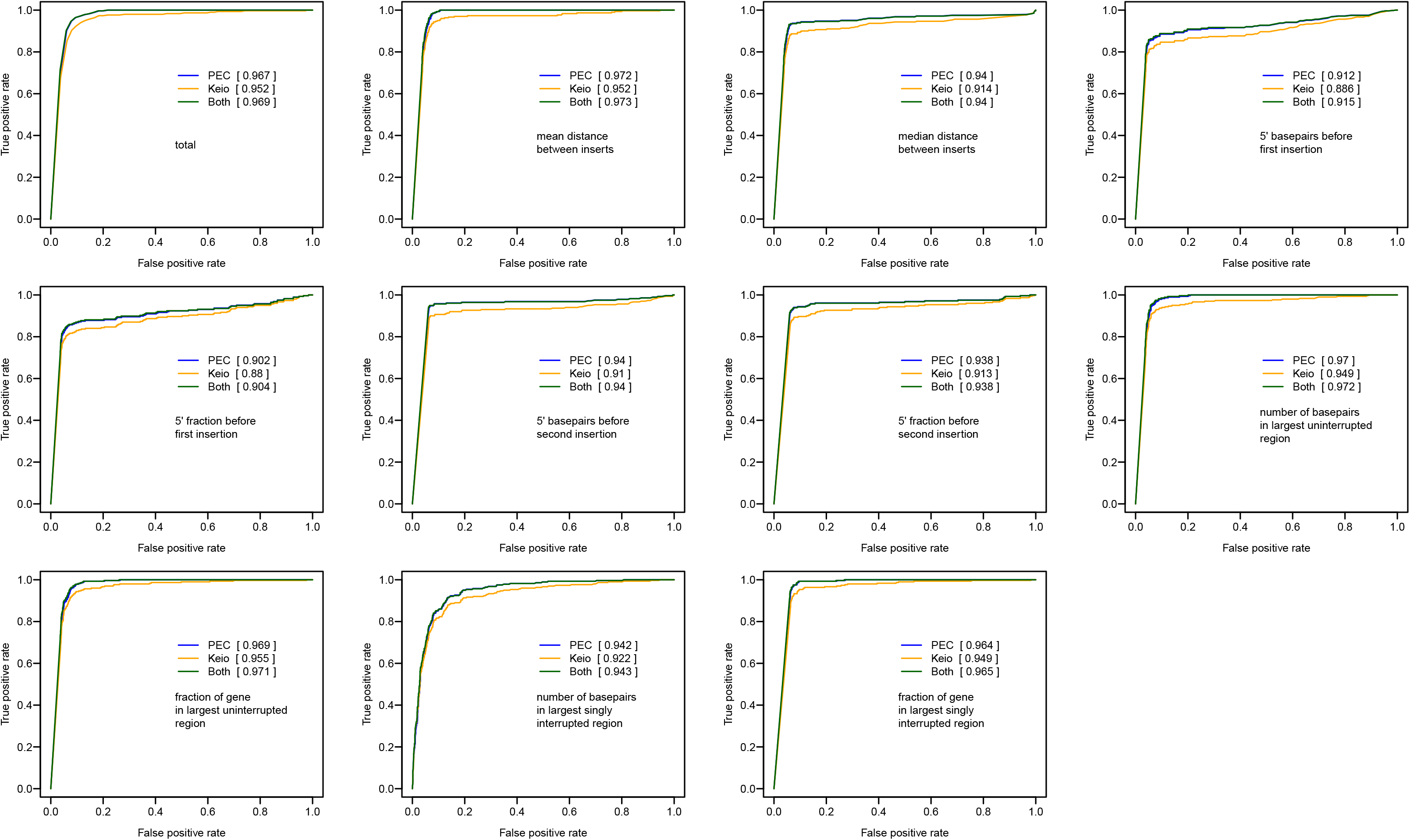
ROC curves showing the predictive power of various features. To select a feature that was the best predictor of essentiality in *E. coli* orthologues, used only ORFs that we had data on essentiality from both the Keio and PEC studies. We then selected transposon insertion patterns that most closely match the essentiality delineations in theses studies. Specifically, we selected the feature that maximized the number of true positive “essential” genes (maximizing the sensitivity) while minimizing the number of FP (maximizing specificity). We selected from eleven (non-independent) features shown here: (1) the total number of insertions; (2) the mean number of bp between insertions; (3) the median number of bp between insertions; (4) the number of bp in the 5’ end preceding the first insertion; (5) the number of bp in the 5’ end preceding the first insertion relative to the total bp in the gene; (6) the number of bp in the 5’ end preceding the second insertion; (7) the number of bp in the 5’ end preceding the second insertion relative to the total bp in the gene; (8) the number of bp in the longest uninterrupted stretch of the gene; (9) the number of bp in the longest uninterrupted stretch of the gene relative to the total length of the gene, and (10) the number of bp in the longest stretch of the gene interrupted by at most one insertion; (11) the number of bp in the longest stretch of the gene interrupted by at most one insertion, relative to the total length of the gene. See the **Methods** section for more details of this analysis.

**S3 Fig.**
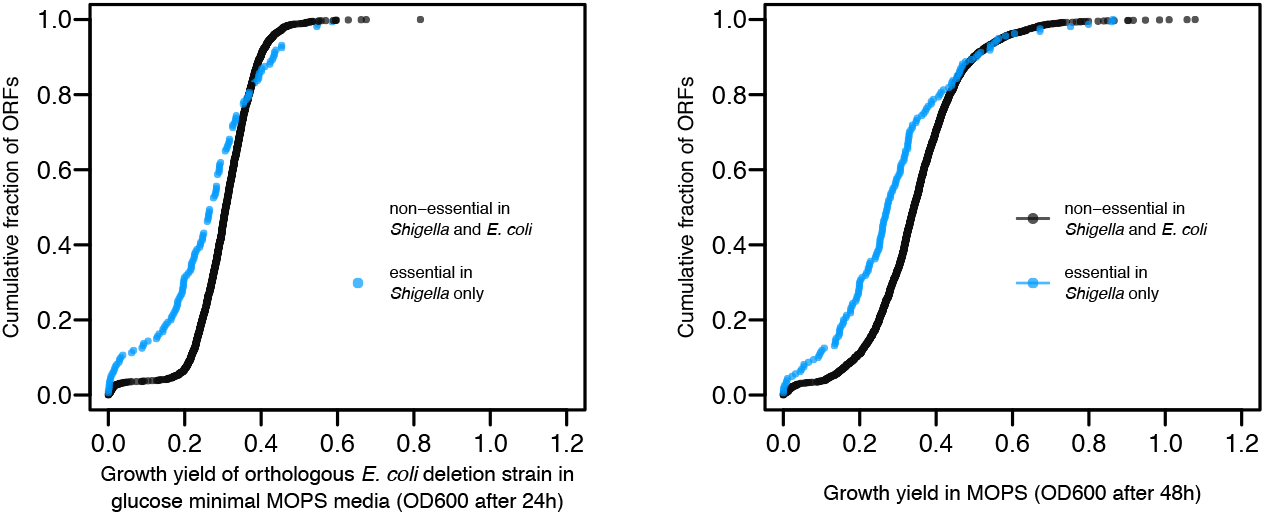
Analogous plots to that shown in Fig. 3, for growth in minimal glucose MOPS media after (A) 24 and (B) 48 hours. In both cases, we find that the shift is less pronounced than that observed for LB.

**S4 Fig.**
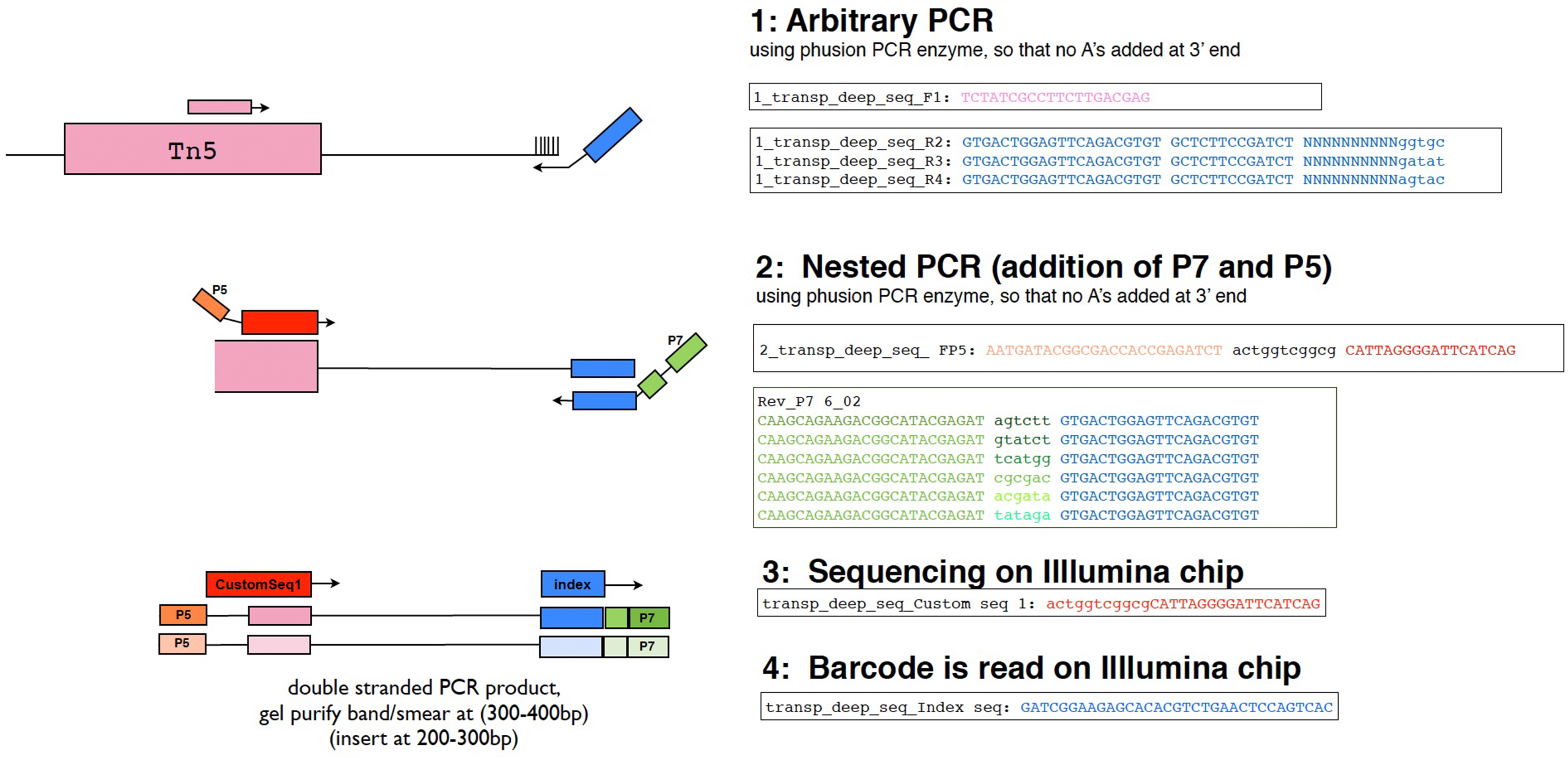
Schematic of the primer positions used for Illumina sequencing of transposon insertions.

**S5 Fig.**
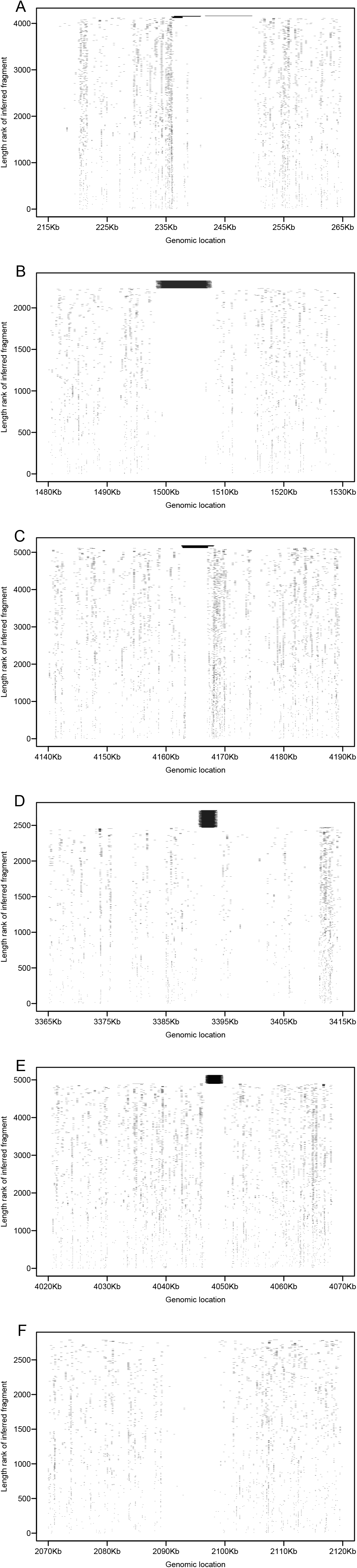
Inferred fragment lengths of perfectly mapped reads across several genomic regions. For each plot, the inferred fragment lengths are arranged by increasing length (ranked on the y-axis). Thus, very long fragments are present at the top of the y-axis. Most fragments have lengths between 100bp and 400bp; a small number have lengths over 1000bp or more. It is very likely that these are not the true insert sizes, but appear that way because of large scale deletions in our *Shigella* clone compared to the clone present in the NCBI genome database; see **Methods** for more details. (A) A region of the chromosome in which a complicated series of rearrangements has occurred, leading to paired end reads perfectly matching to different locations in this region. 45 mapped read pairs span more than 1.5 Kbp, a size that is not concordant with the majority of insert sizes. (B) A genomic region where an approximately 10Kbp deletion occurred, removing a region containing the *yeaKLMNOP* operon. 92 mapped read pairs span more than 8.5Kbp. This region is flanked by two IS elements. (C) A region where an approximately 4Kbp deletion occurred, removing two genes with no *E. coli* K12 orthologues. 68 mapped read pairs span more than 4 Kbp, and again this region is flanked by two IS elements. (D) A genomic region where an approximately 2Kbp deletion occurred, removing *yhdW*. 244 mapped read pairs span more than 2Kbp, and the region is flanked by two IS elements. (E) A deletion in the region of the chromosome containing *S4145 (yiaN)*. 232 mapped read pairs spanned more than 1.8 Kbp, and this region is also flanked by two IS elements. (F) A region of the chromosome containing the *rfb* operon. Most of the genes within this operon are uninterrupted by transposons. However, we find no evidence that this is due to a deletion of this region in our *Shigella* clone, as we find no reads mapping across the region; a small number of reads map within the region; and the closest IS elements are 15 Kb upstream of *rfbJ* and 20 Kb downstream of *rfbA*. The genes in this operon have no orthologues in *E. coli* K12.

**S6 Fig.**
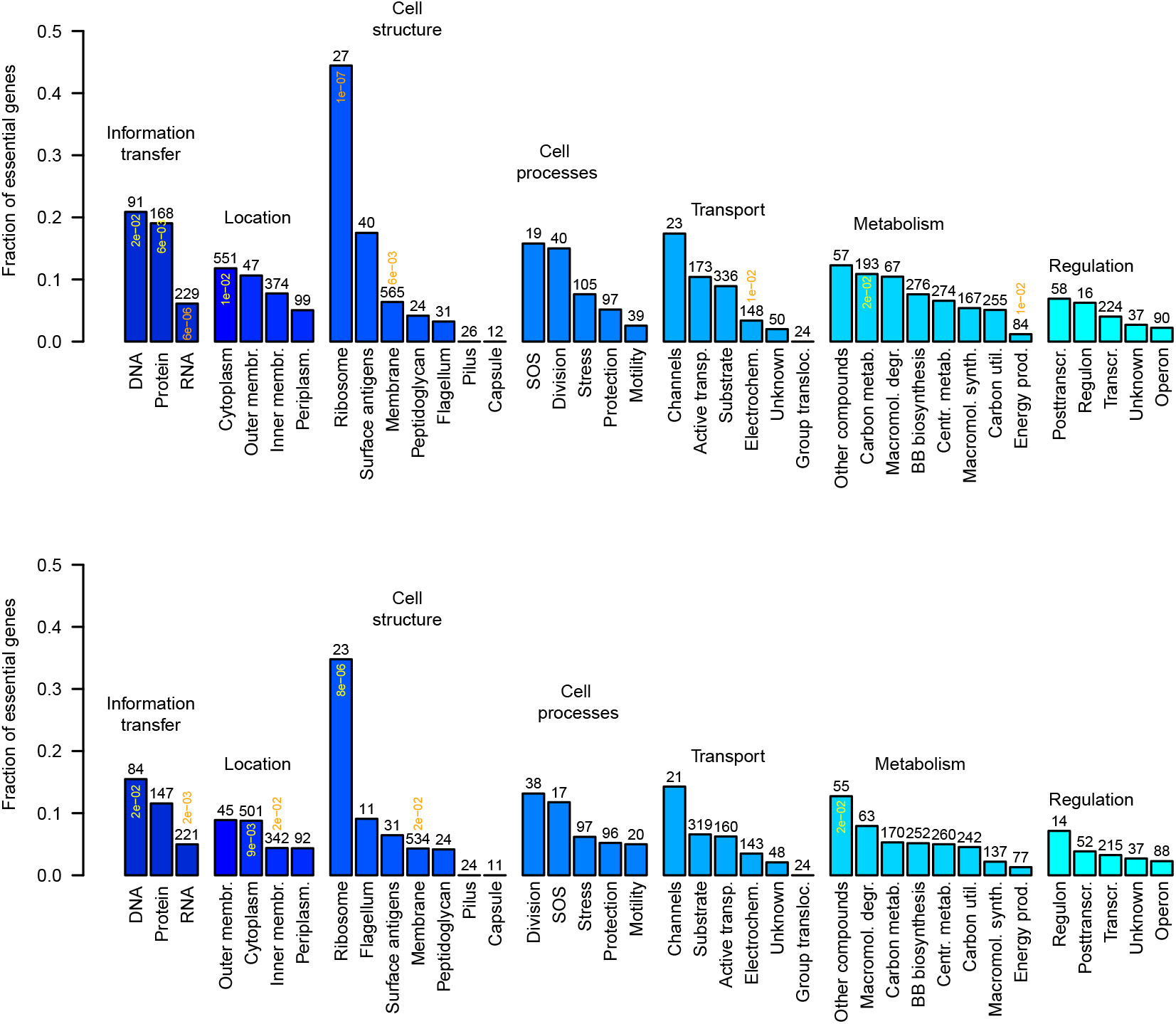
Functional categories of non-essential *E. coli* genes that are enriched (or depleted) for essential *Shigella* genes. We classified genes according to function using the MultiFun classification system [63]. For this analysis we considered only genes that are non-essential in *E. coli*. We find that genes uniquely essential in *Shigella* are enriched in some functional categories. For example, of the 27 ribosomal proteins identified as non-essential in *E. coli*, we identify approximately 45% as being essential in *Shigella*. In contrast, of the 565 membrane proteins identified as non-essential in *E. coli*, we find that less than 10% are essential in *Shigella*. Thus, ORFs uniquely essential in *Shigella* are far more likely to function in the ribosome than one would expect. The number of non-essential *E. coli* genes is indicated above each bar; the probability of finding the level of enrichment (or depletion) that we observe in each secondary category is indicated for cases in which this probability is less than 0.025, using a Fisher exact test. (A) All non-essential genes in *E. coli*. (B) An identical analysis excluding all non-essential genes in *E. coli* that exhibit very low growth yields (OD600 less than 0.5 after 22 hours of growth LB). In both cases, the only subcategory notably enriched for essential genes is that containing ribosomal proteins. The only categories appreciably depleted for genes with essential function are genes with function in RNA processing and to some extent, energy production.

**S7 Fig.**
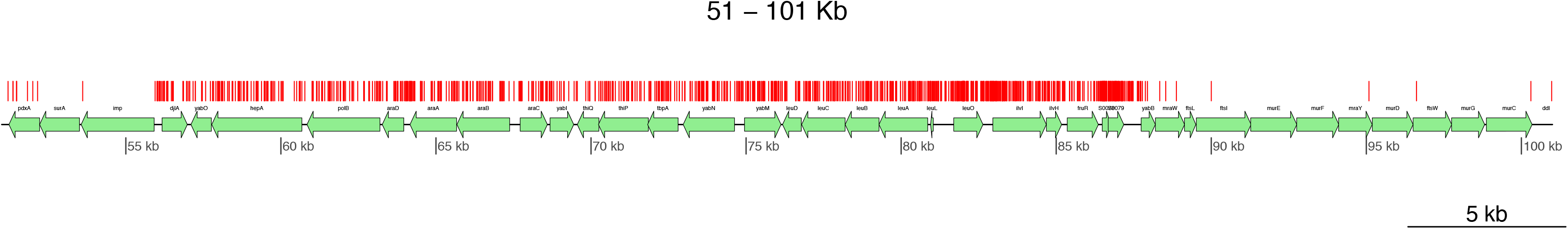
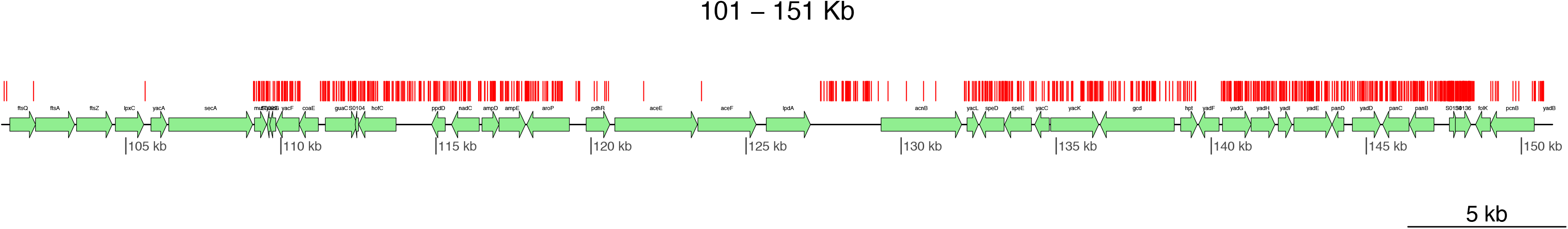
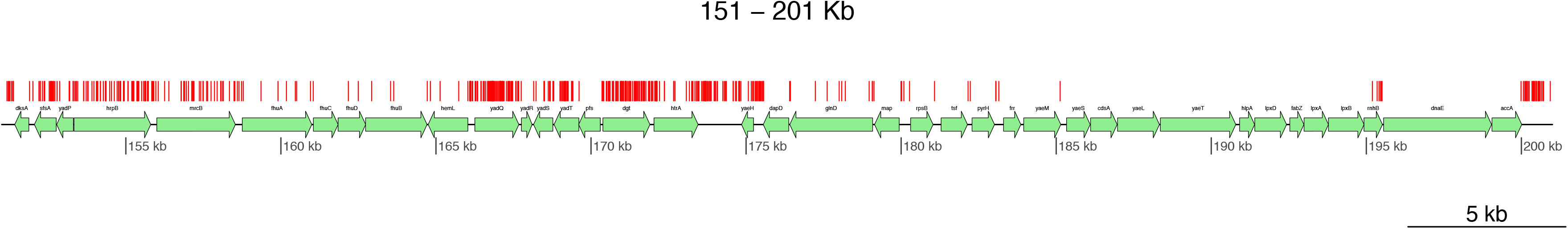
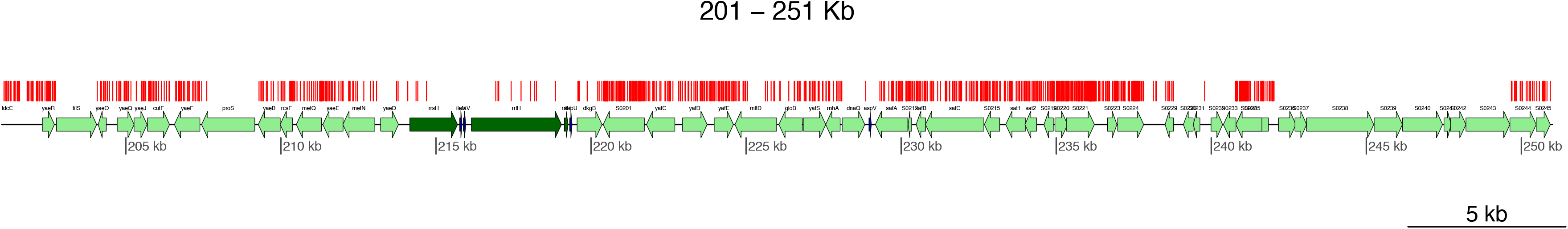
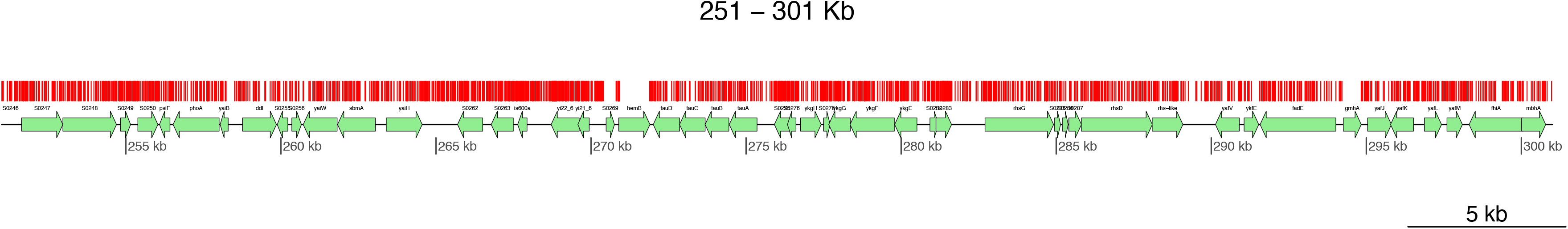
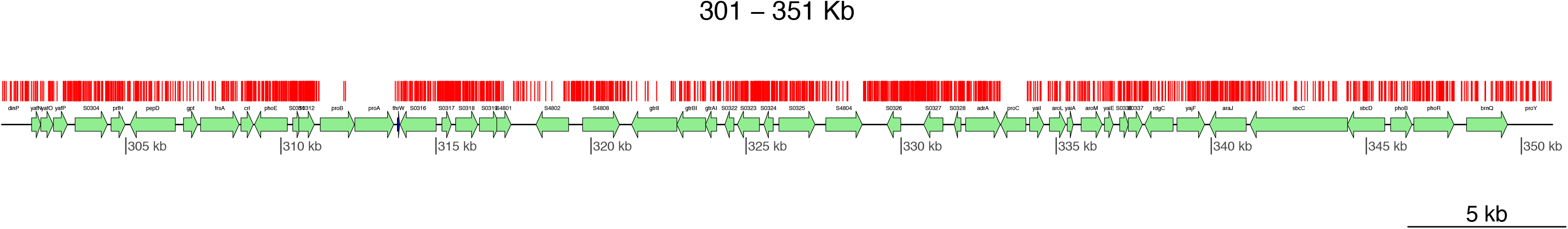
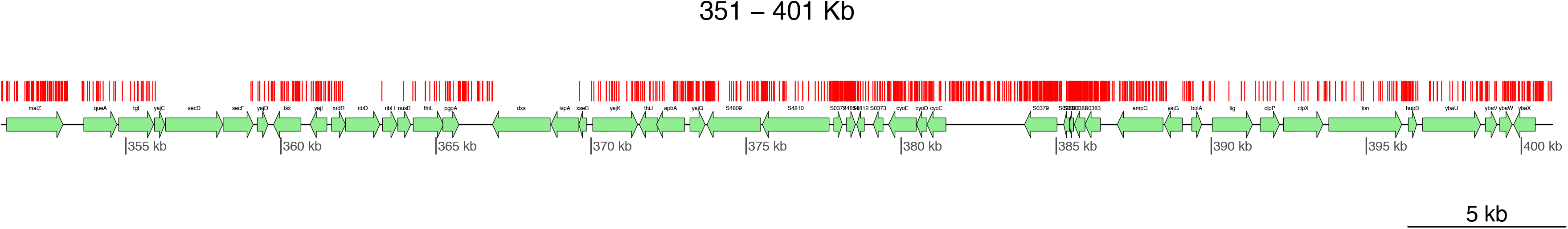
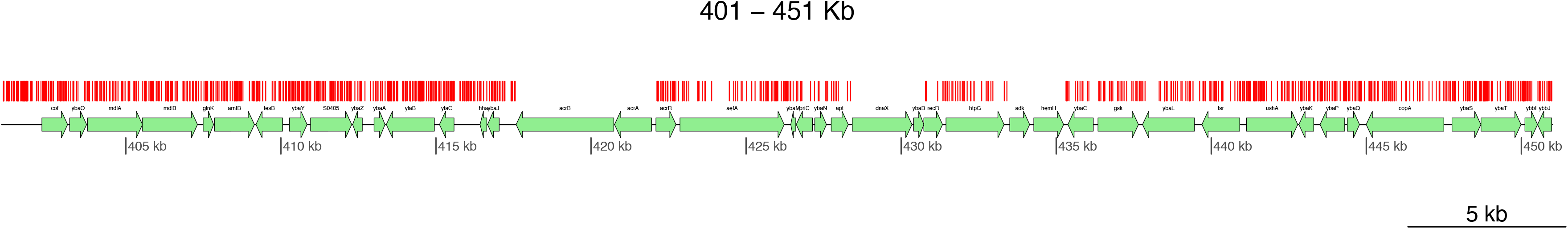
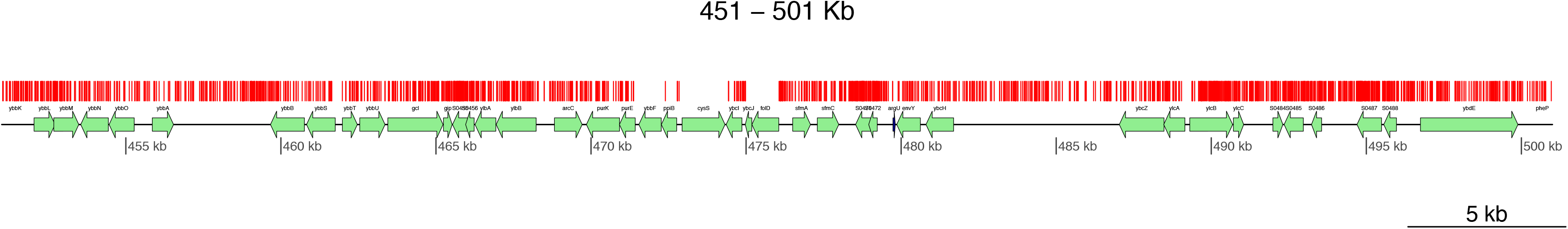
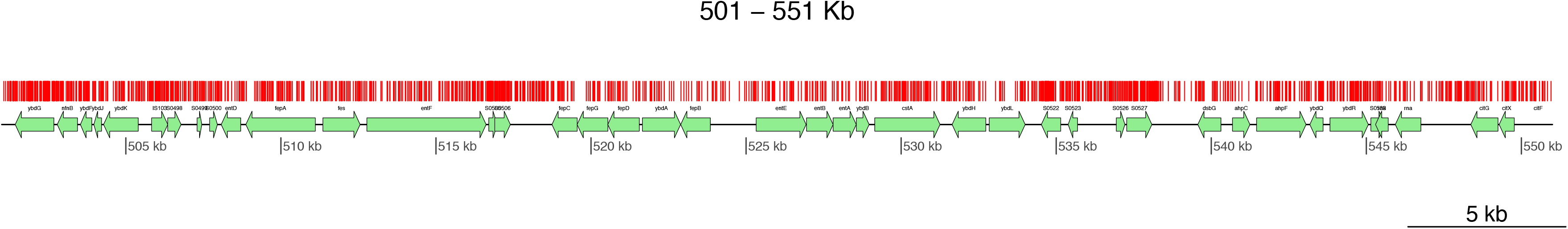
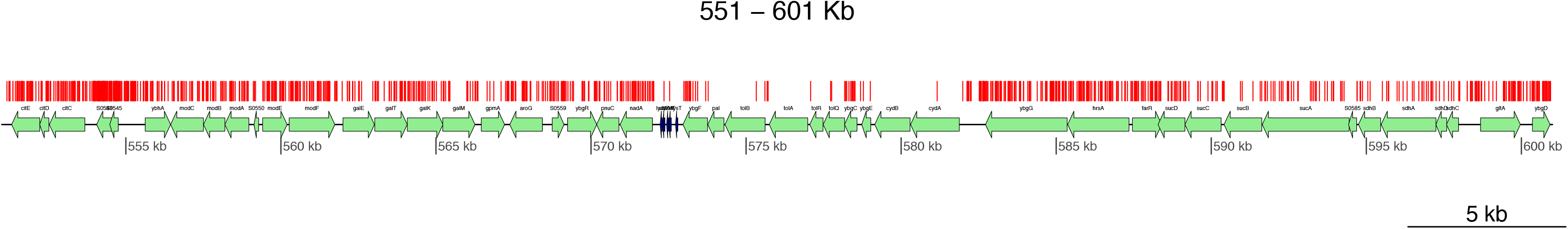
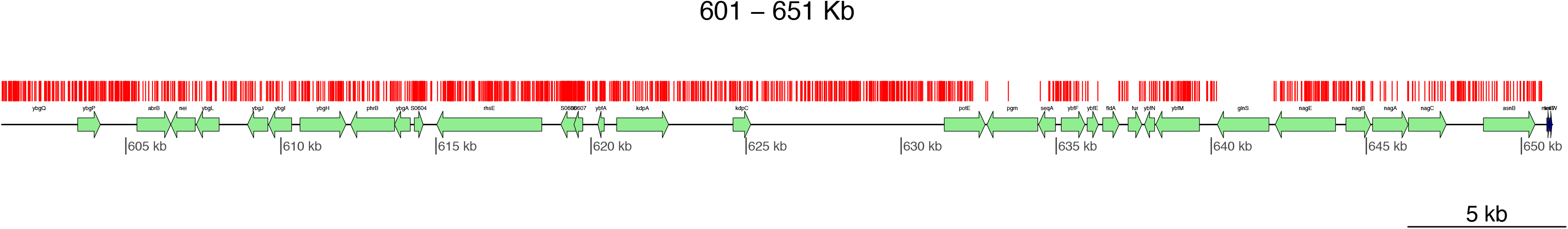
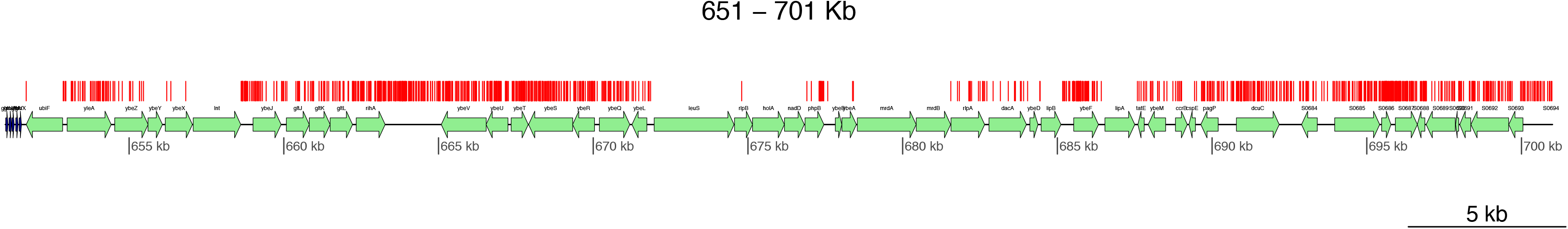
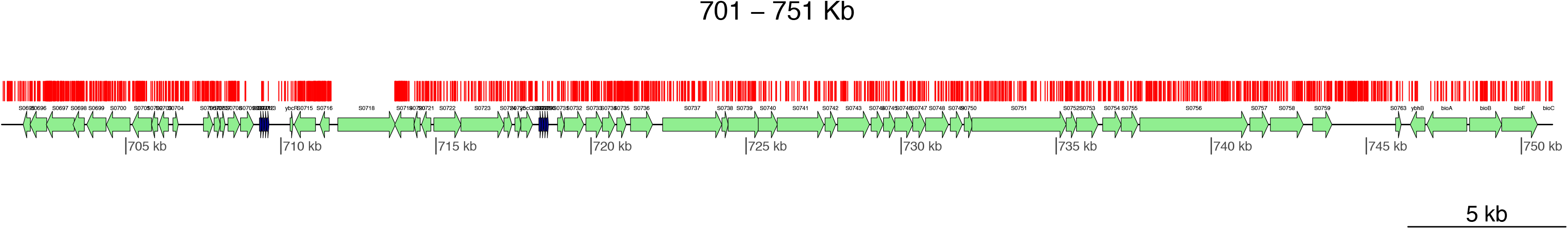
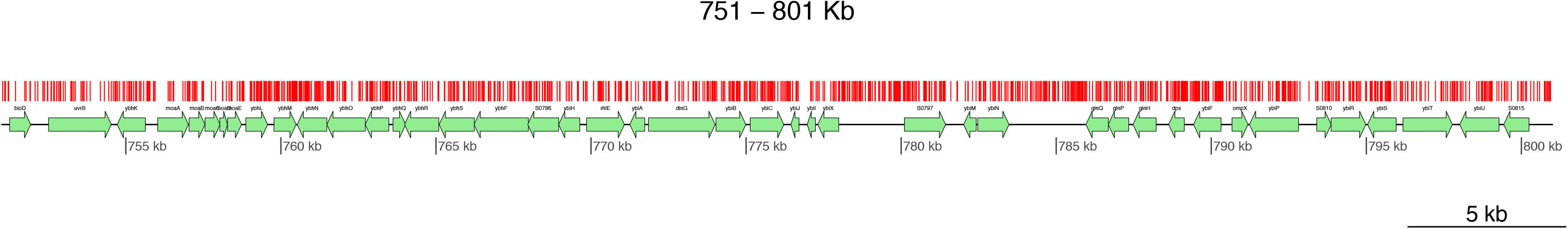
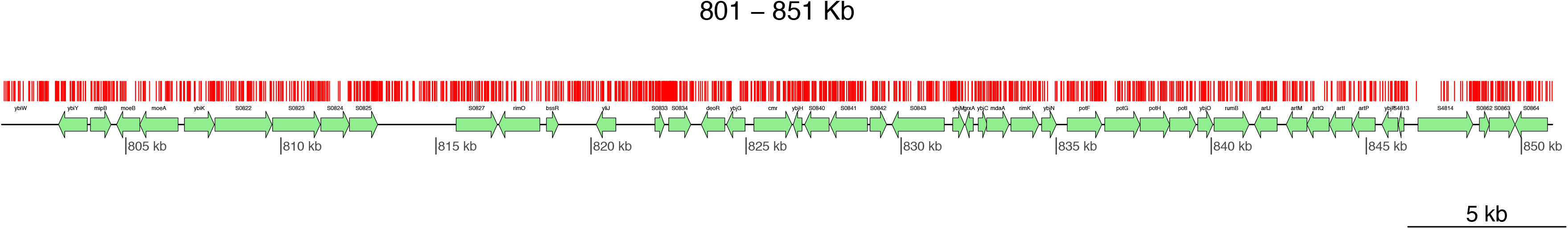
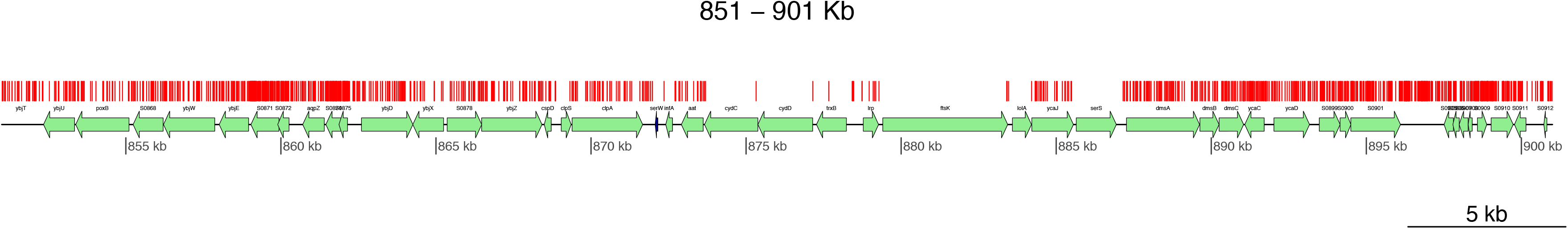
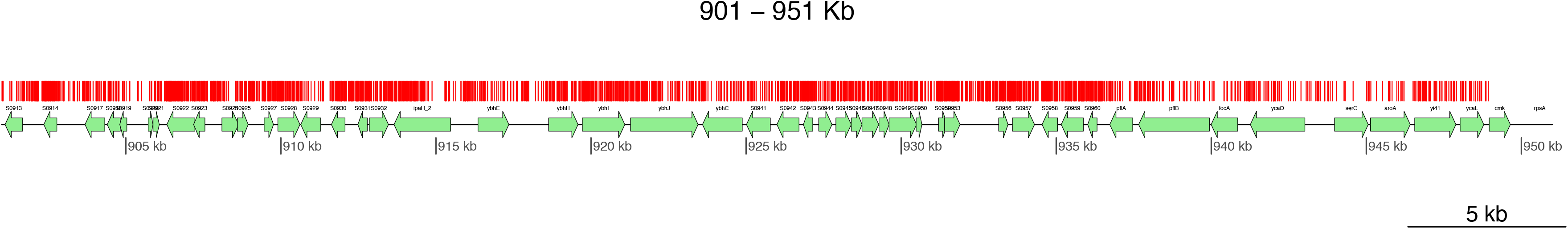
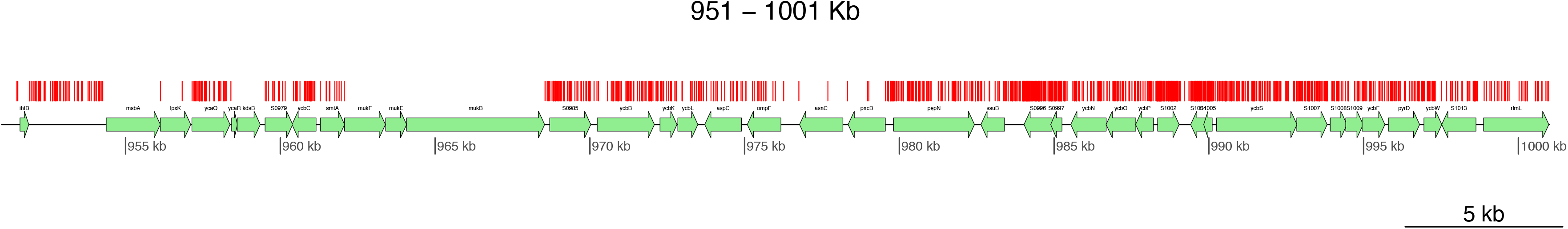
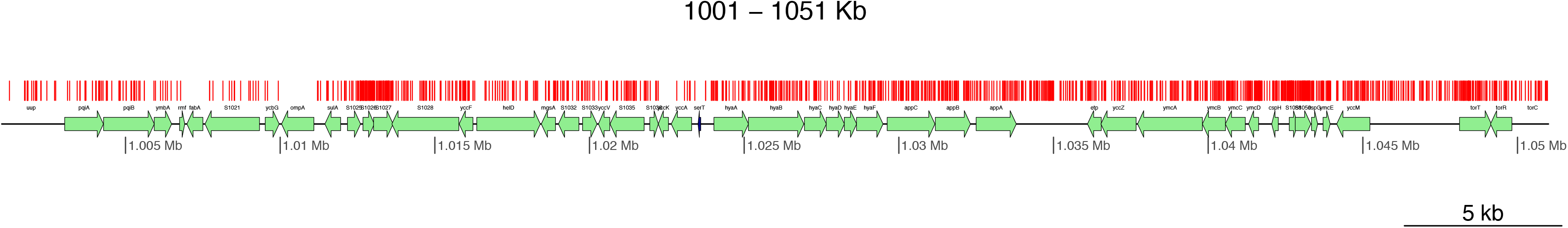
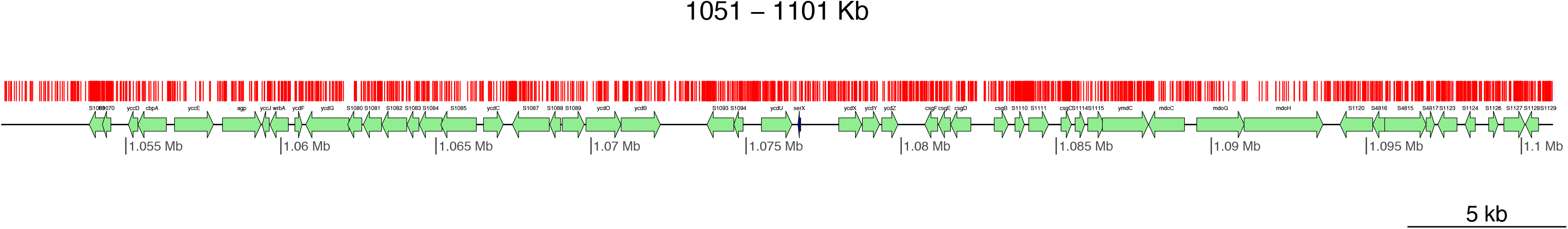
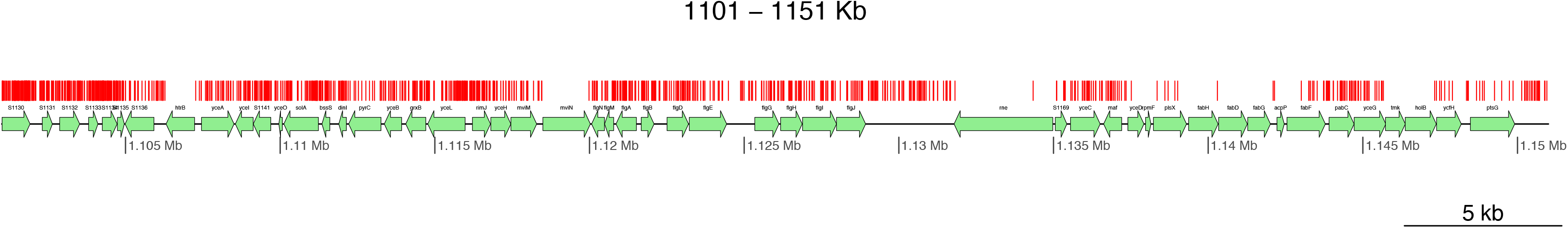
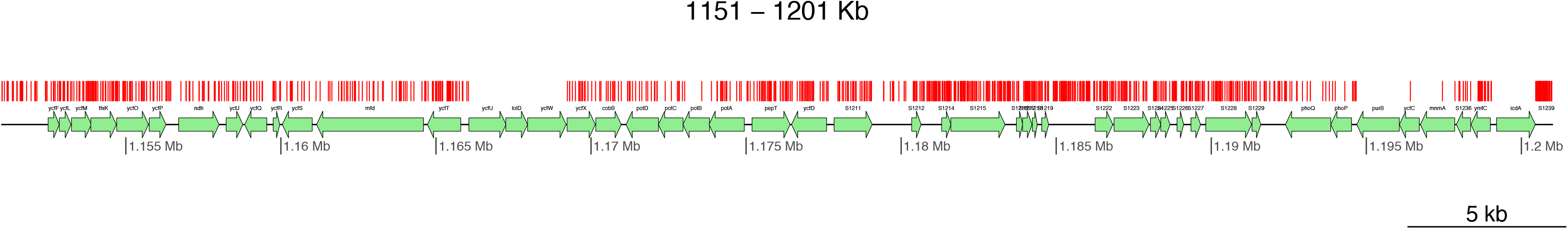
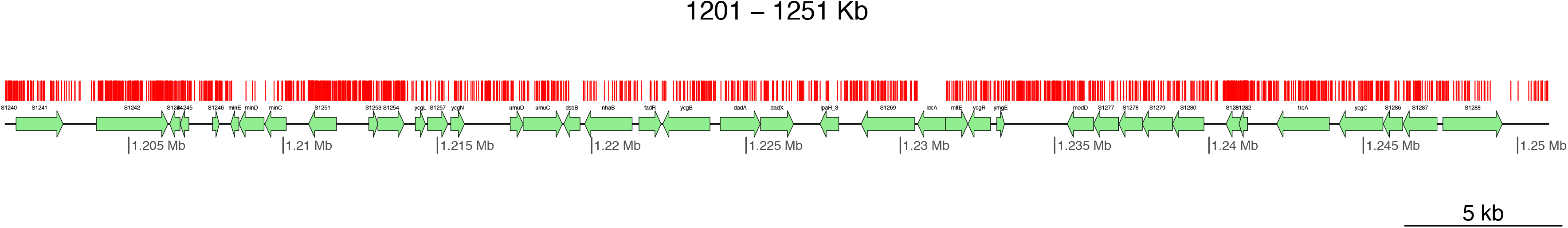
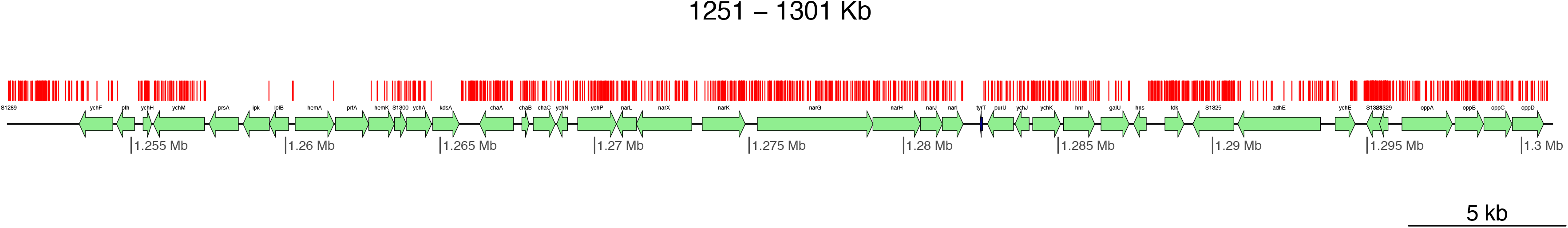
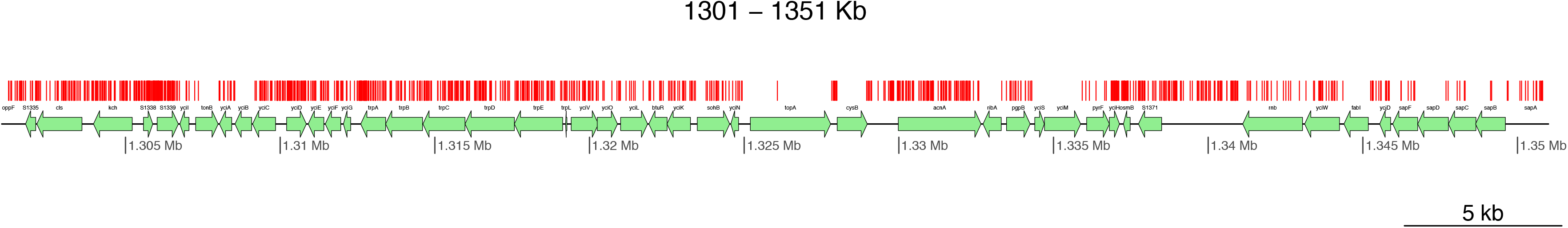
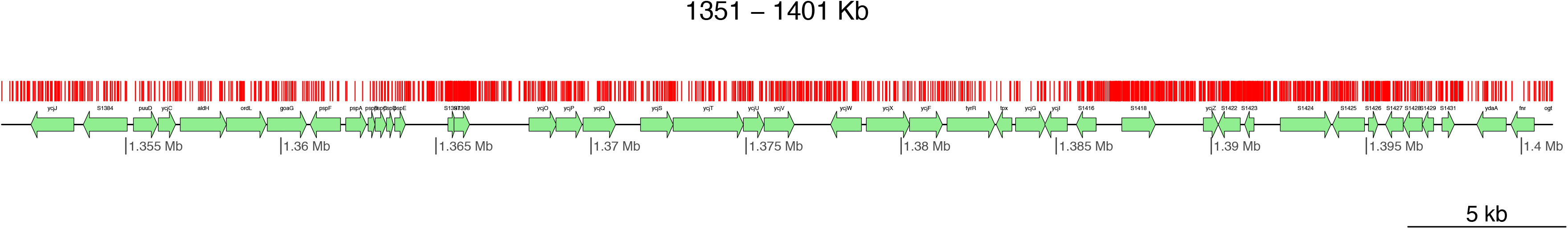
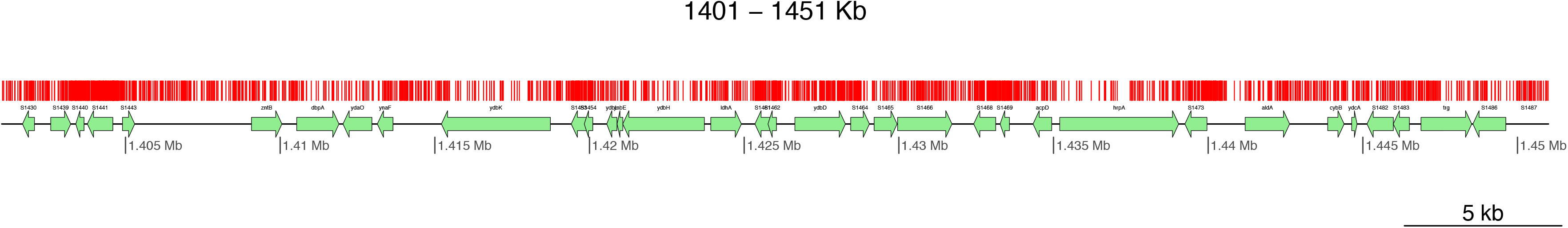
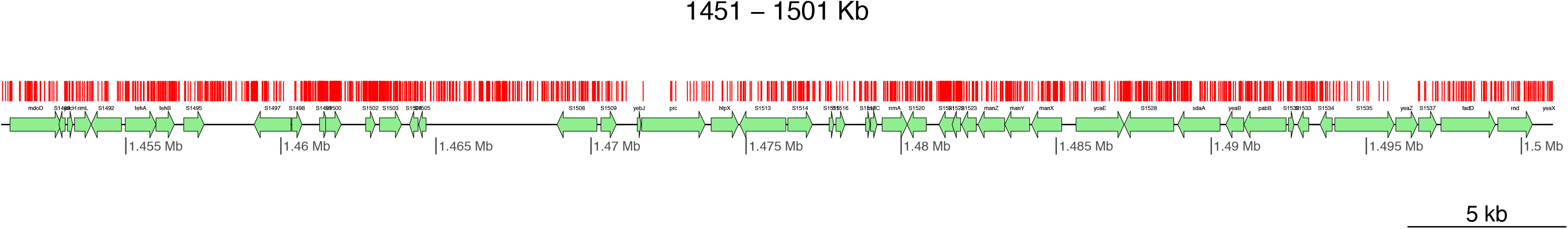
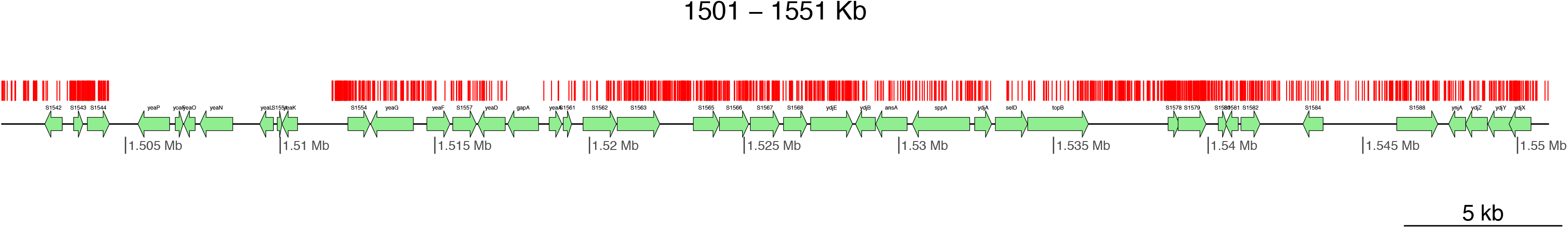
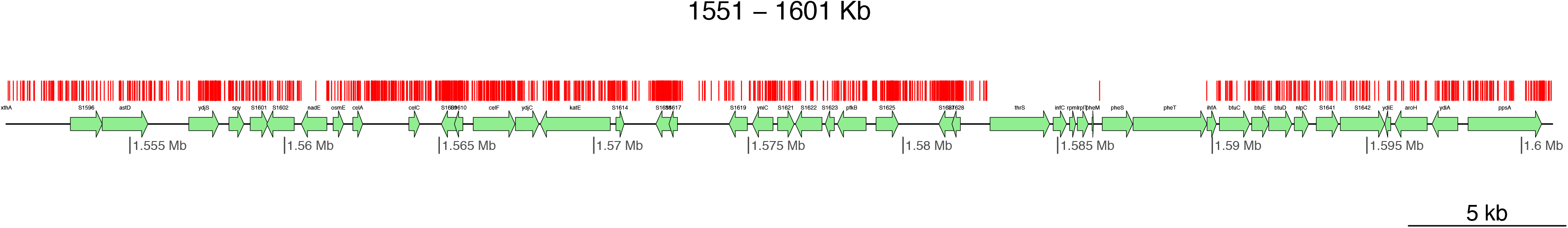
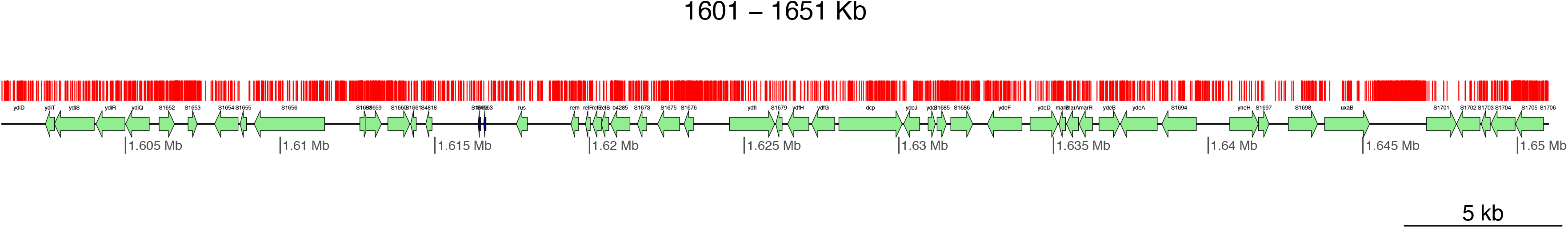
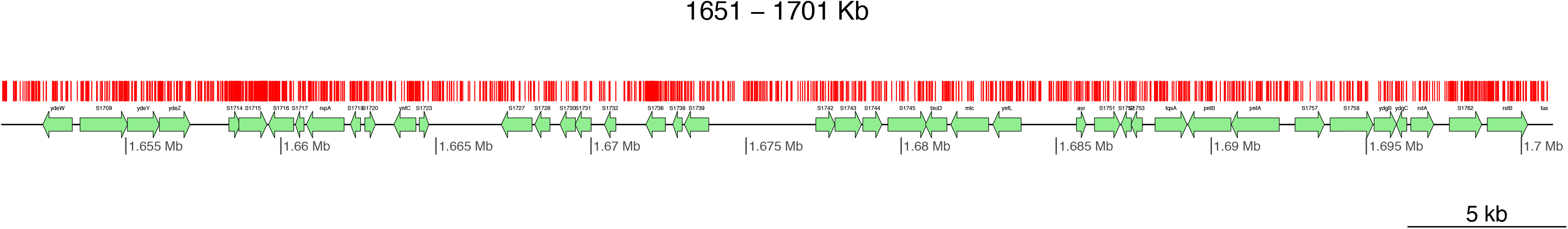
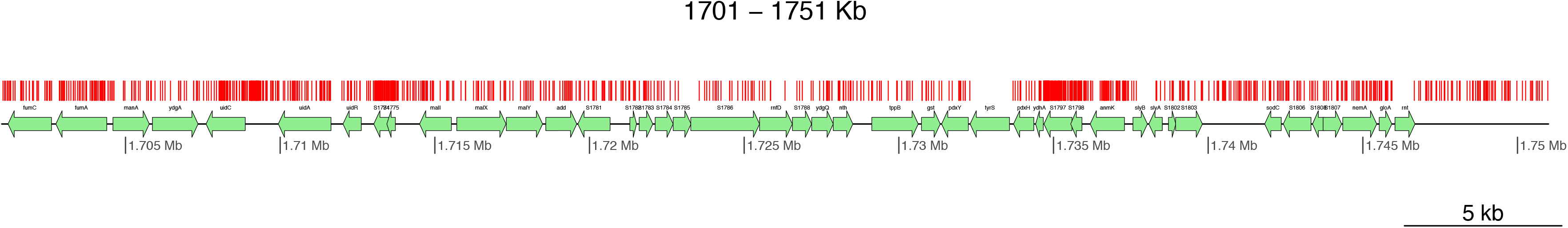
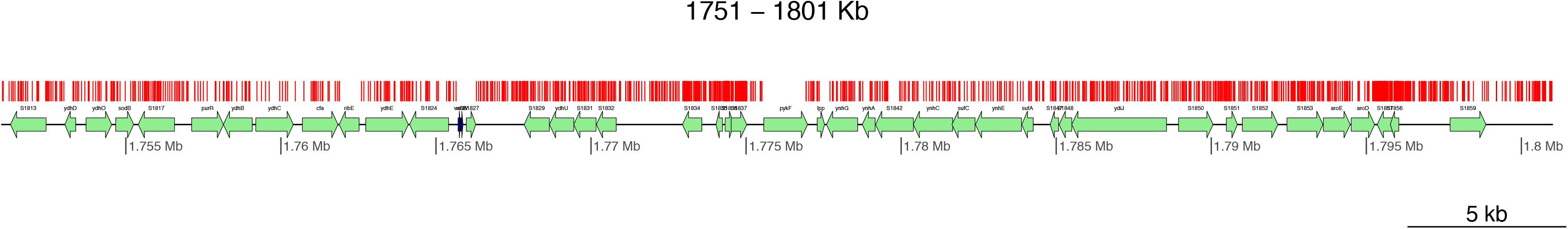
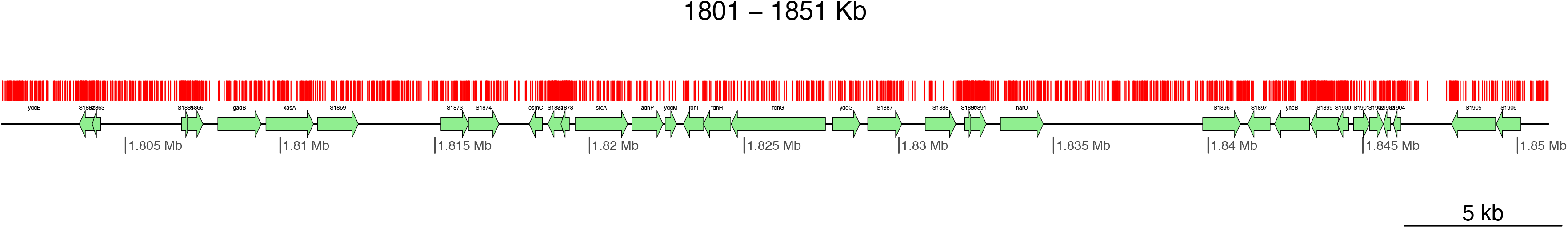
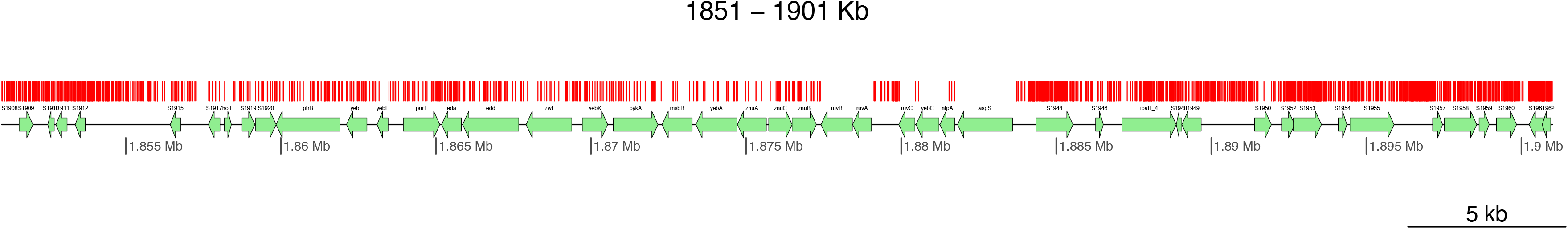
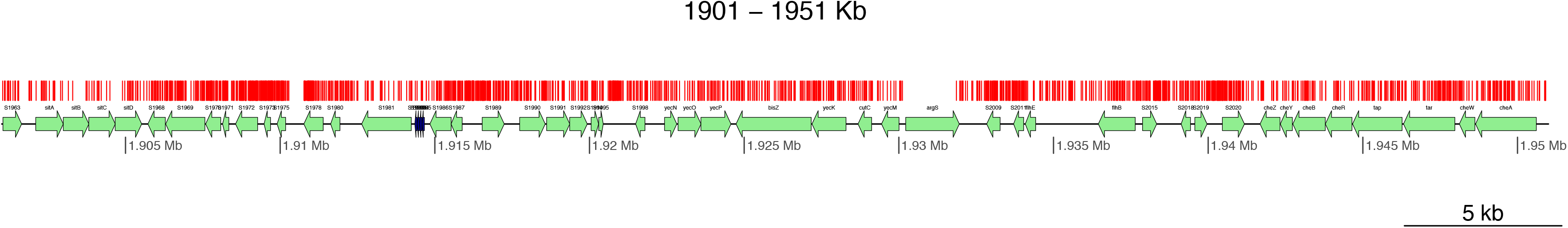
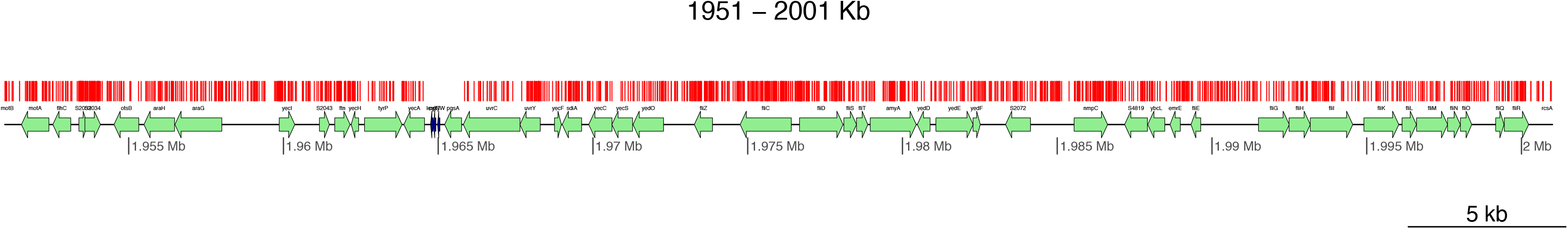
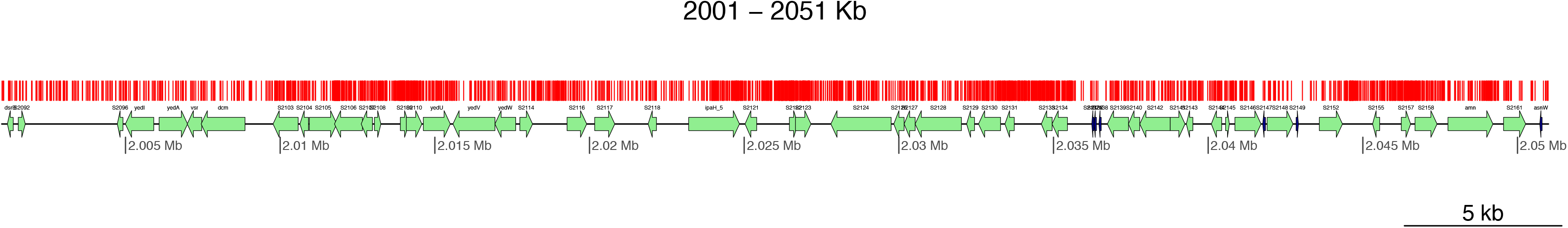
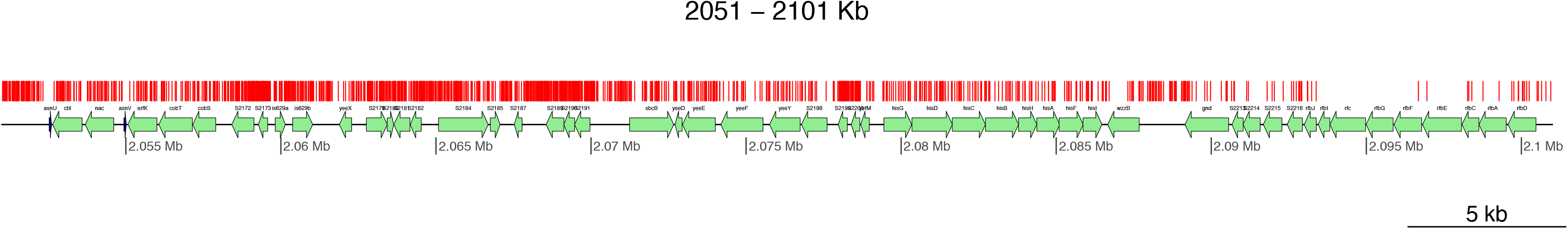
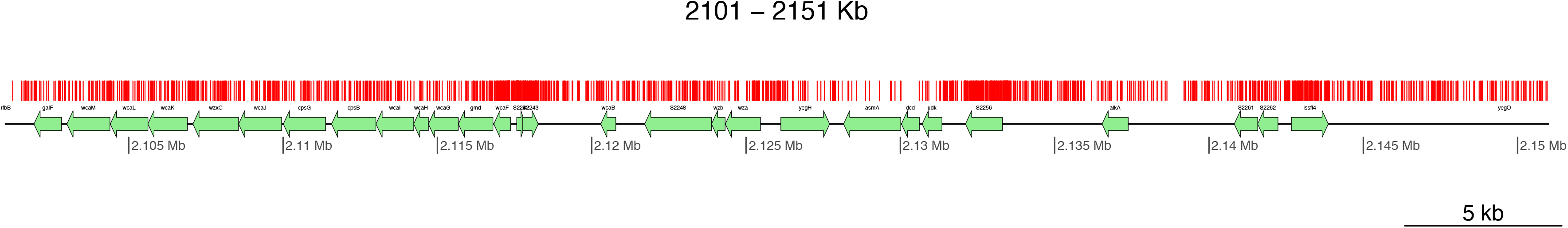
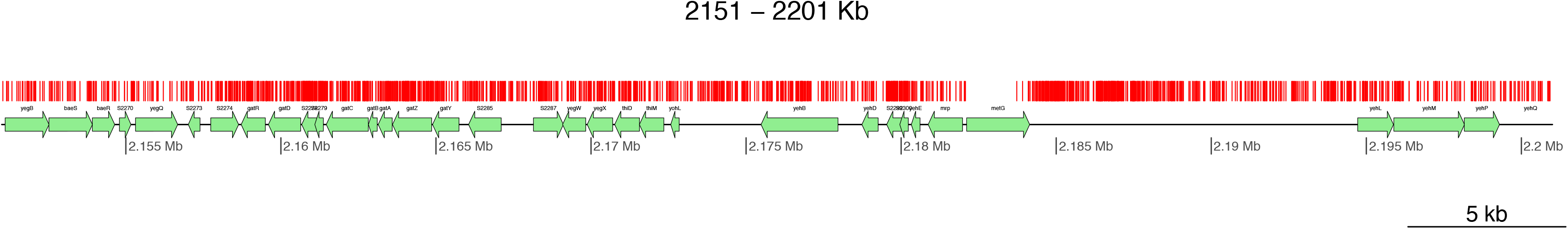
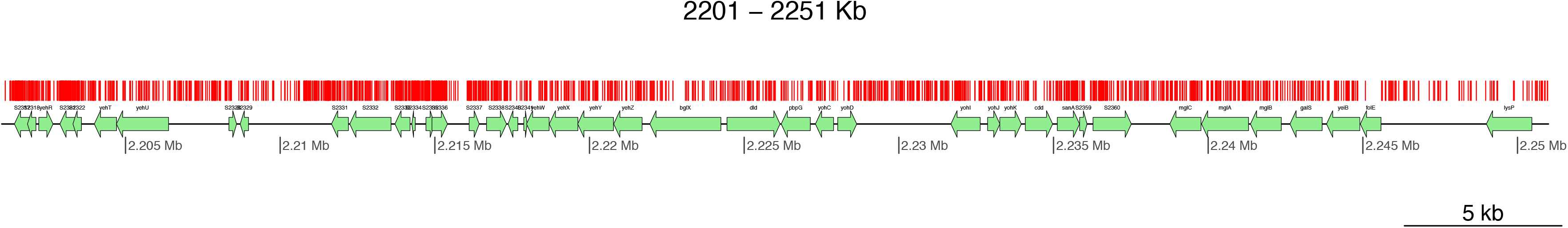
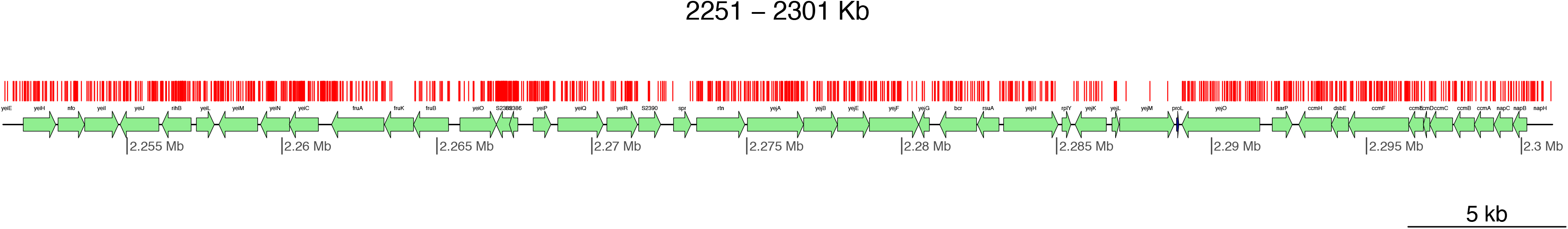
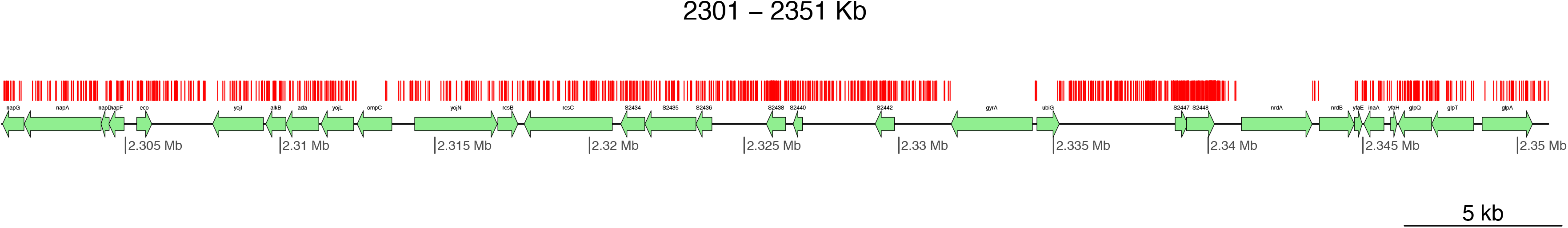
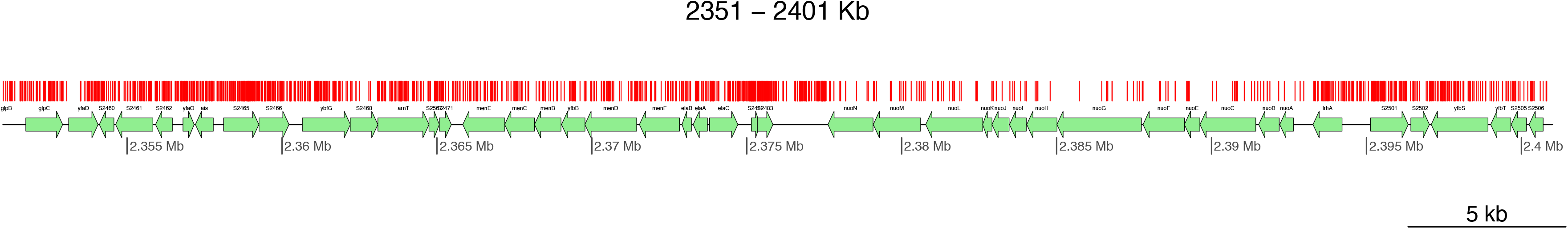
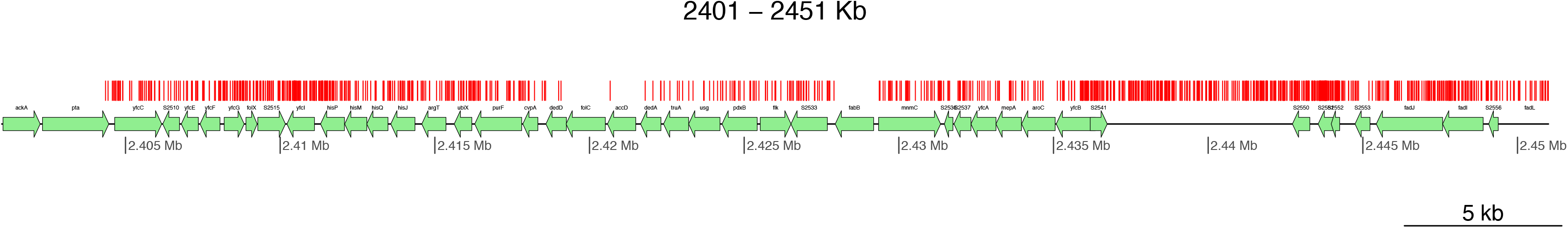
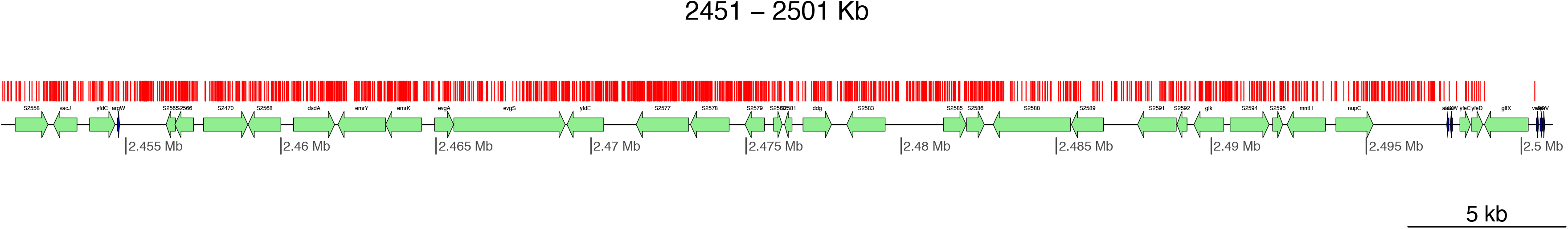
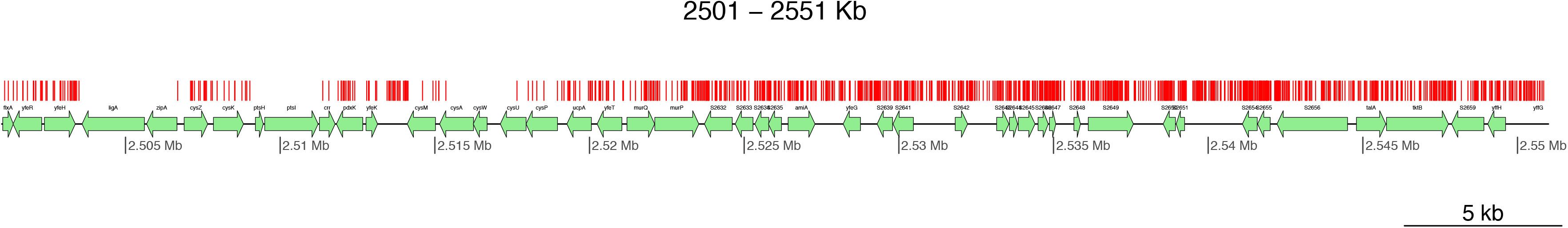
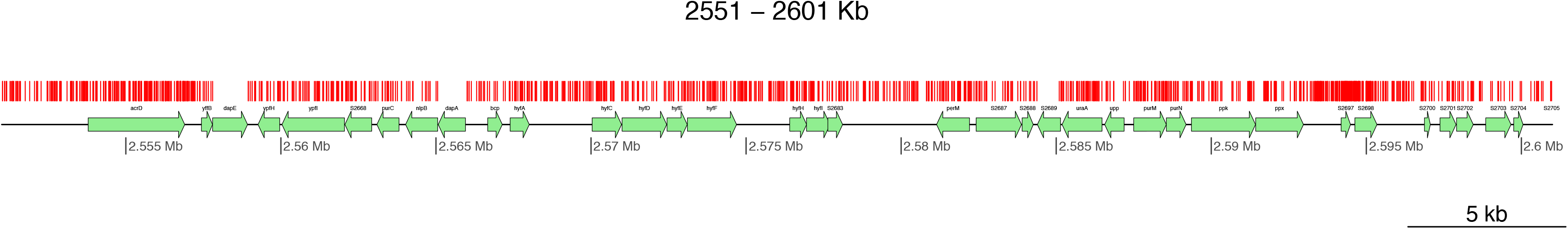
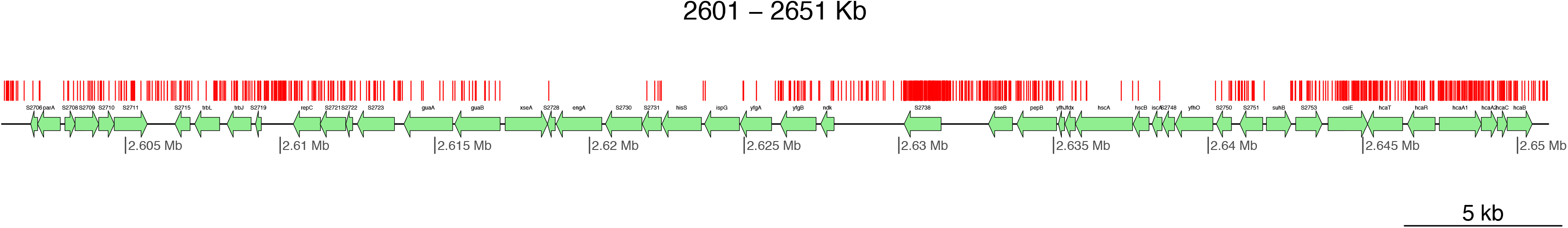
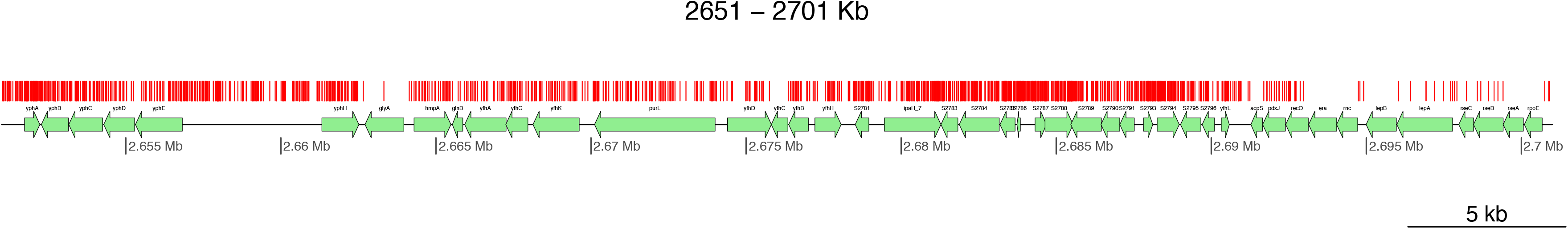
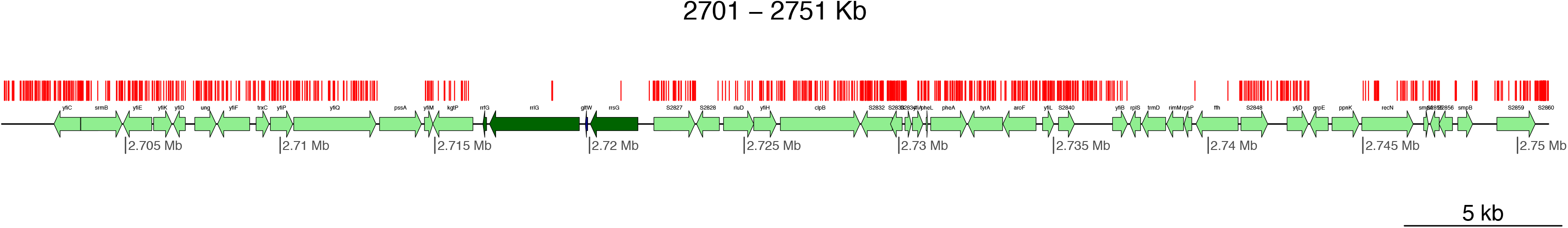
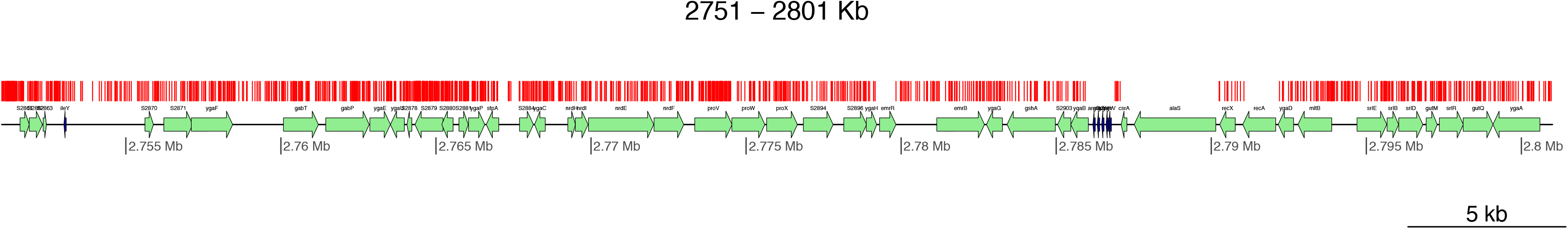
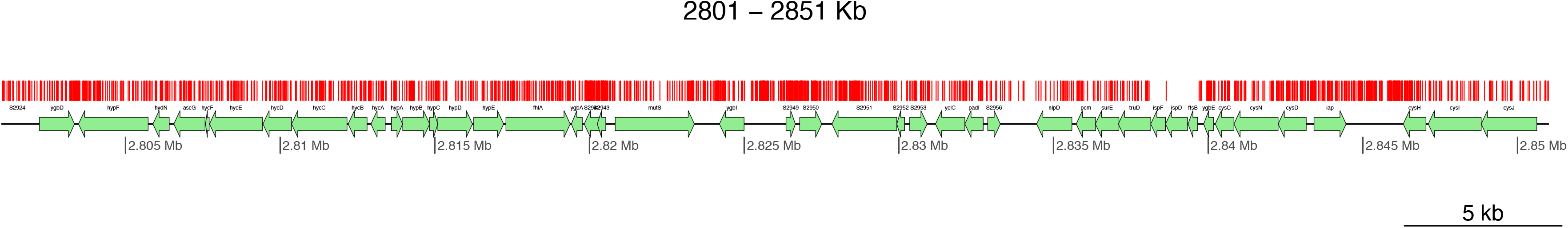
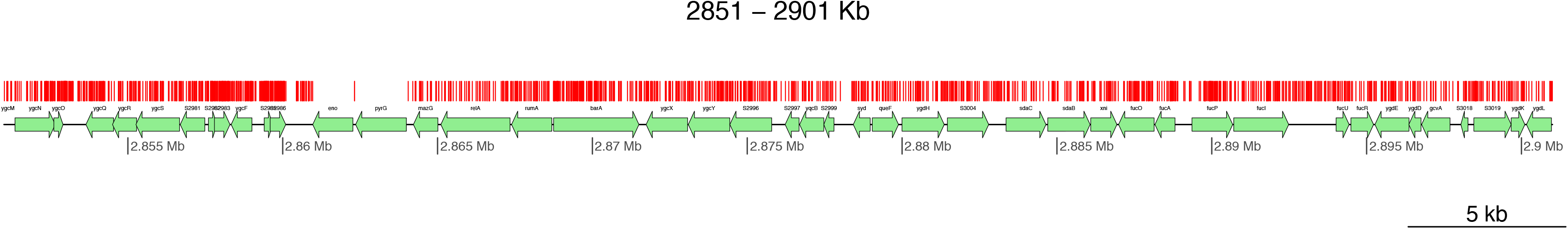
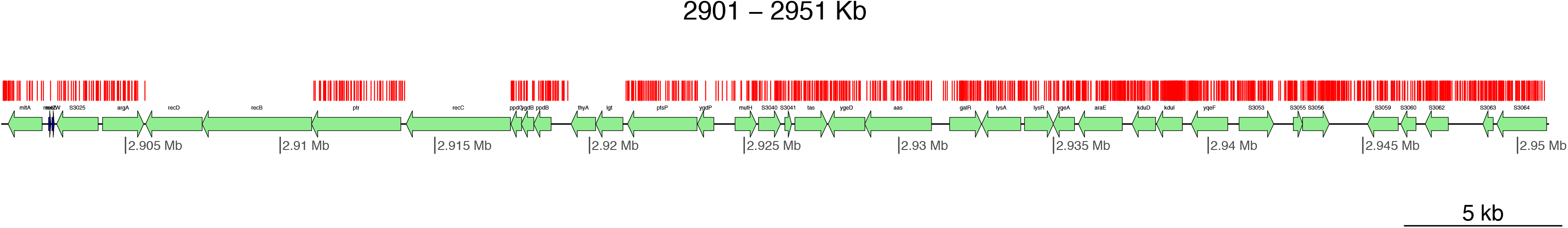
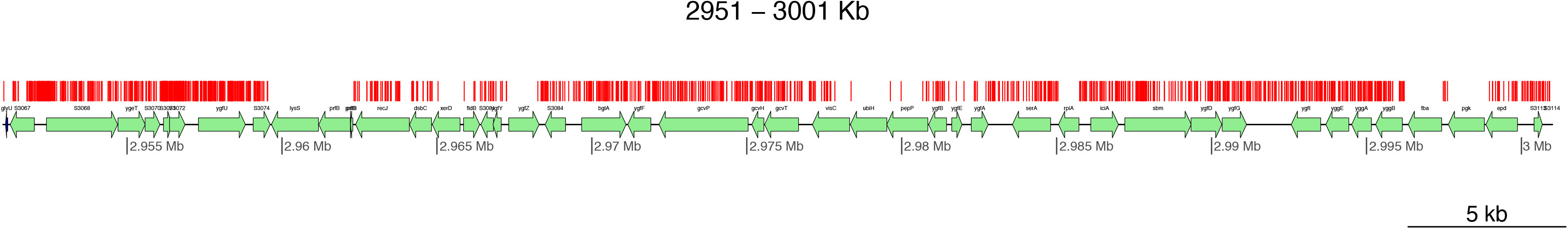
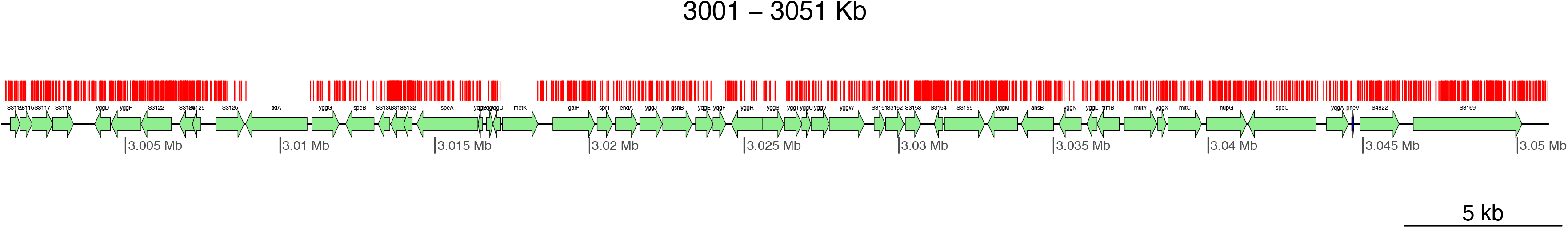
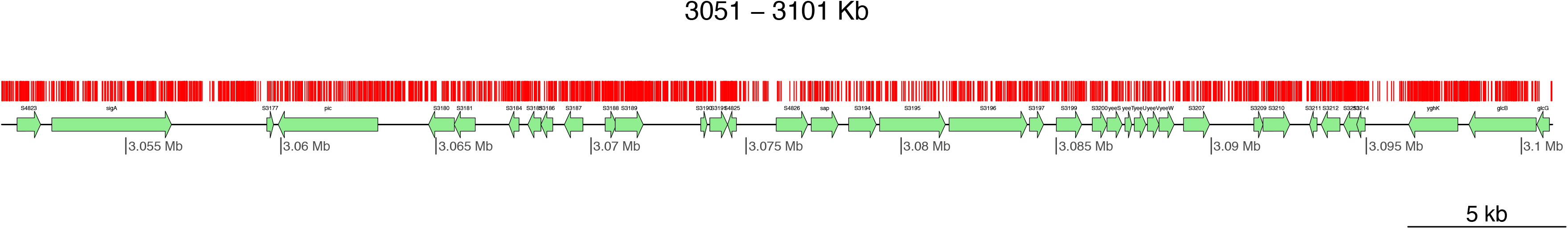
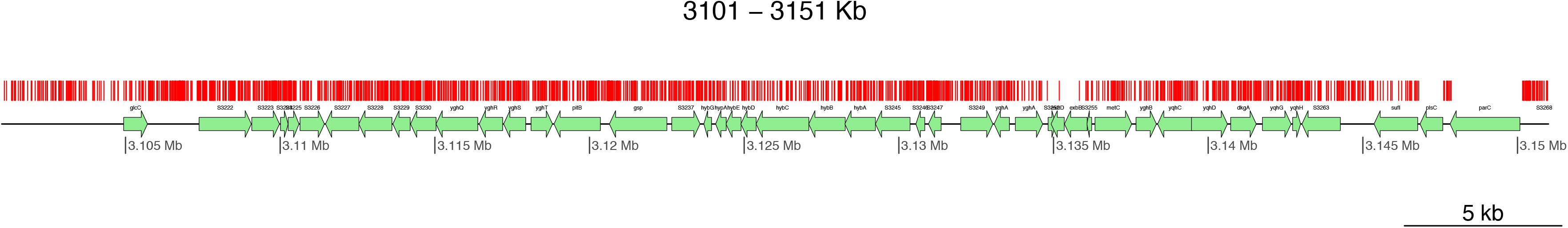
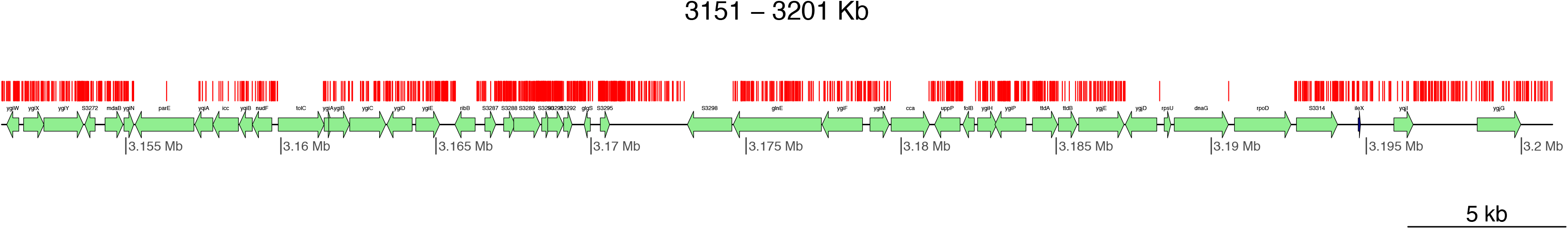
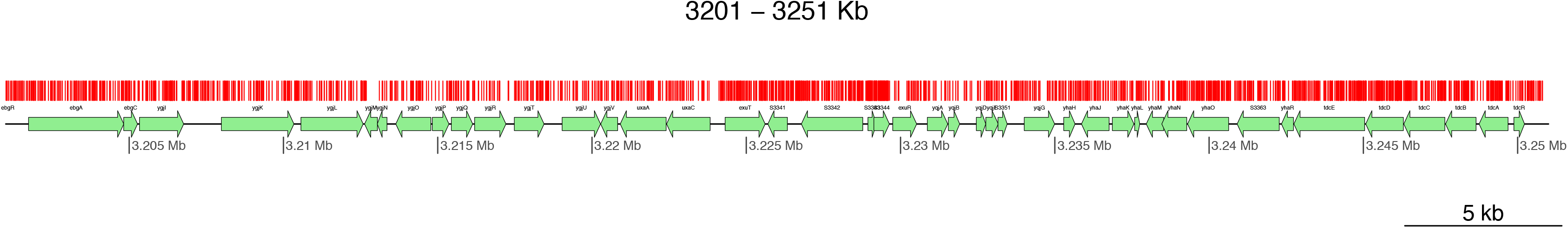
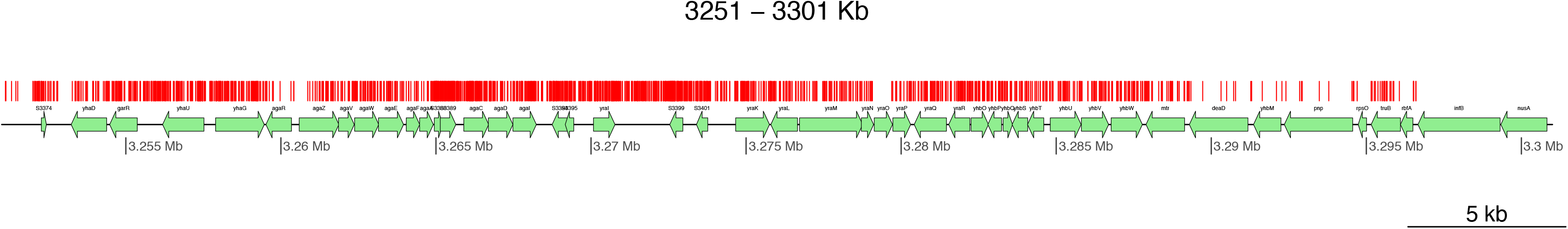
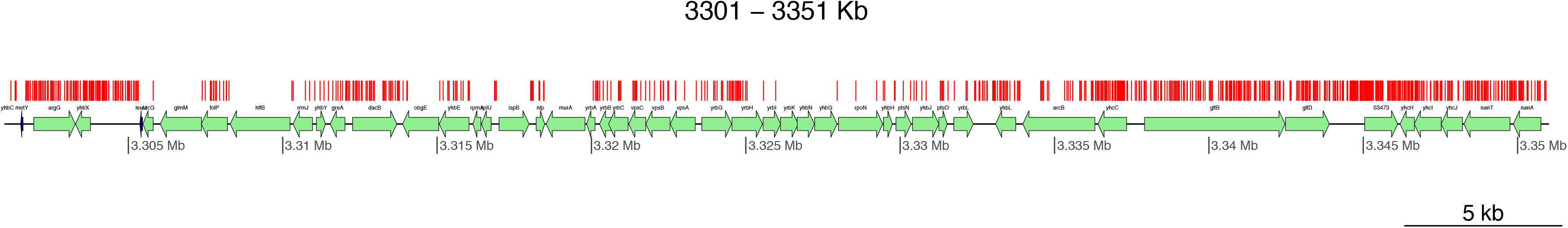
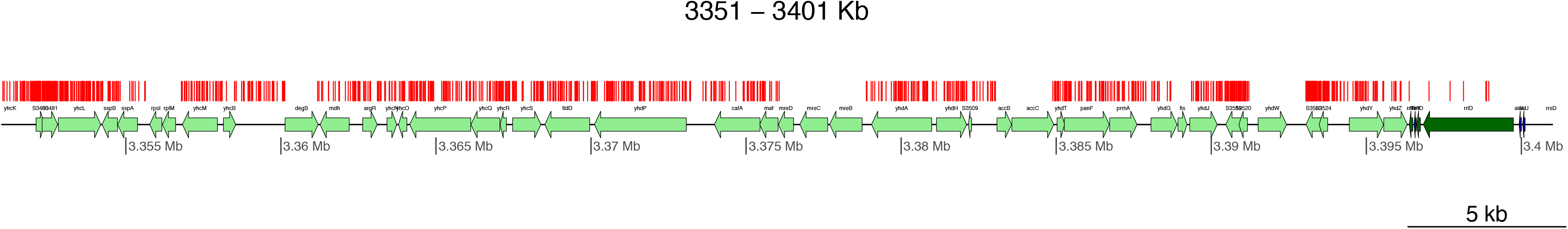
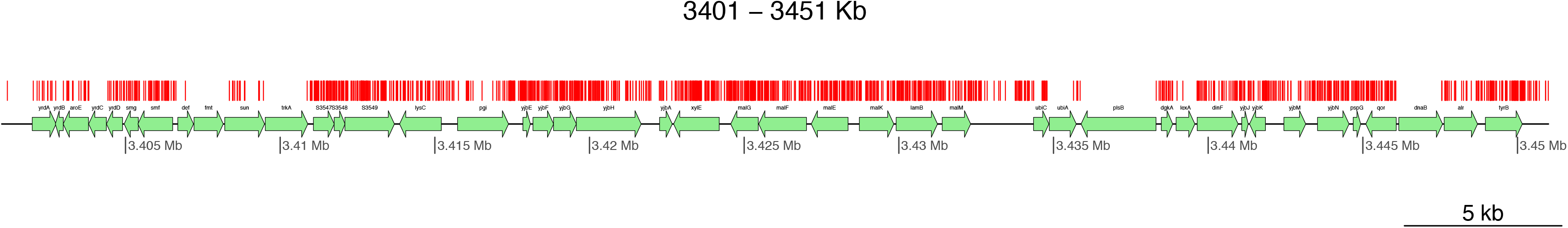
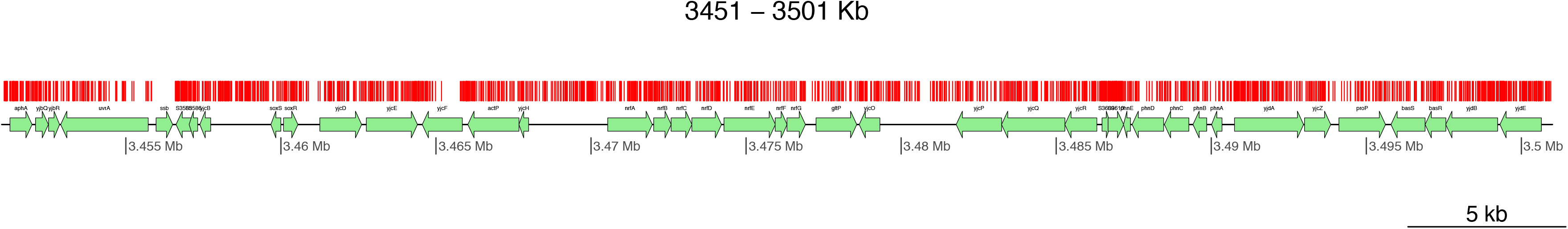
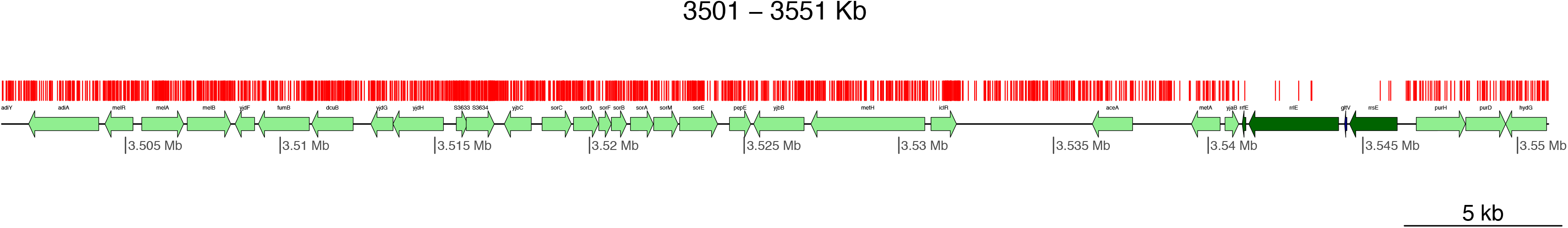
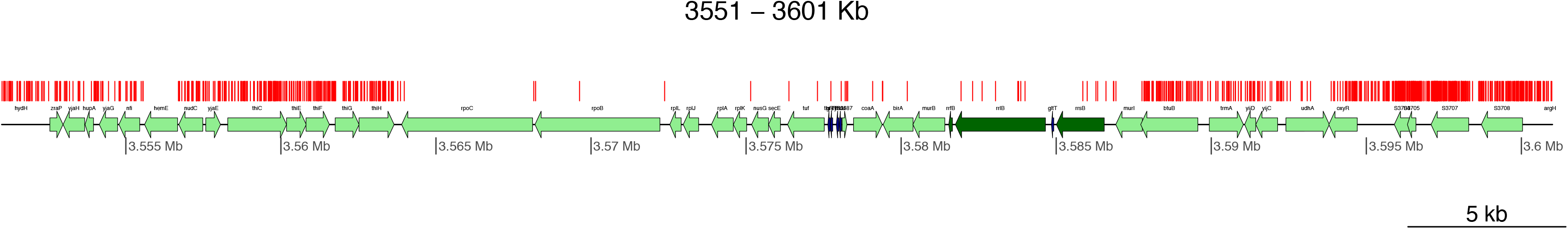
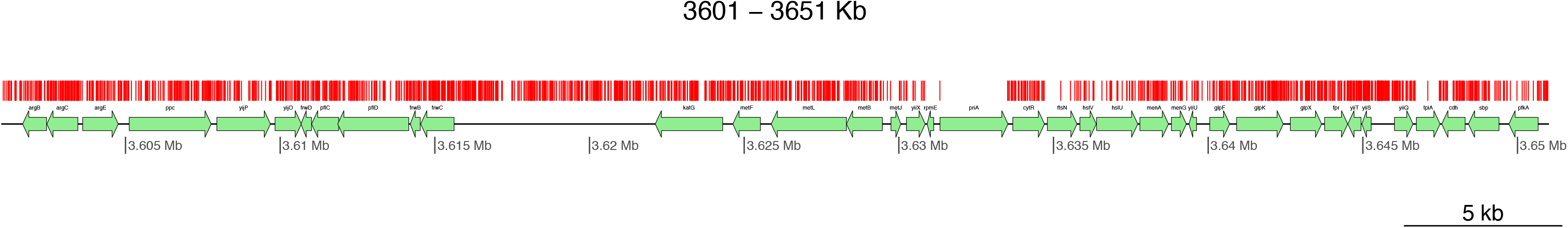
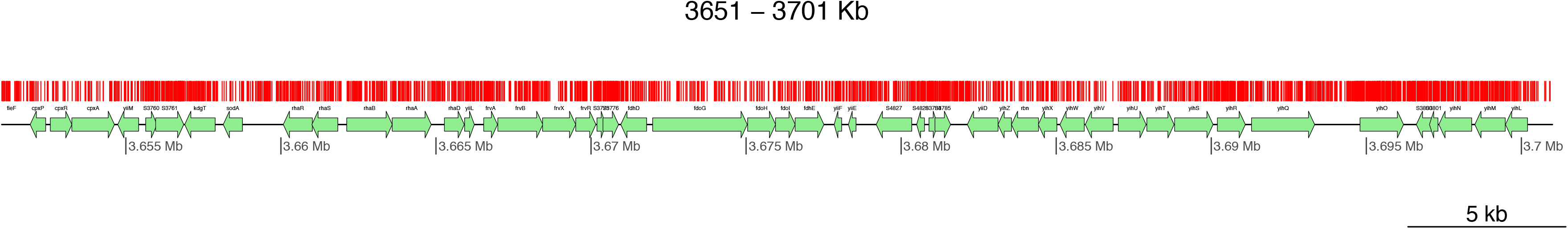
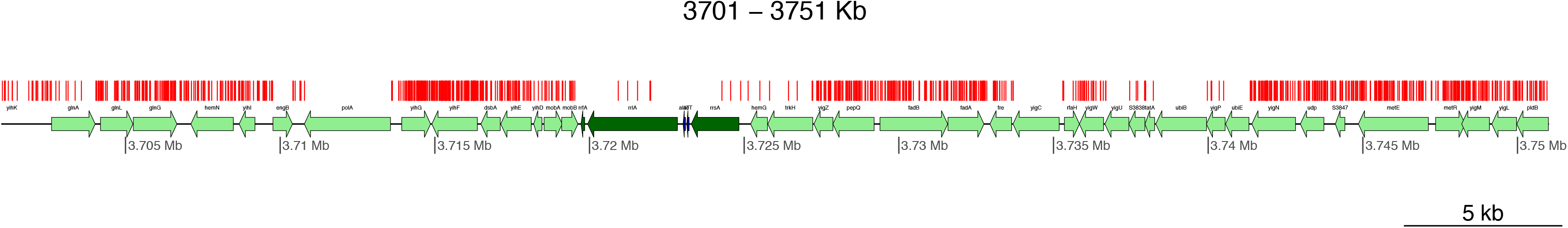
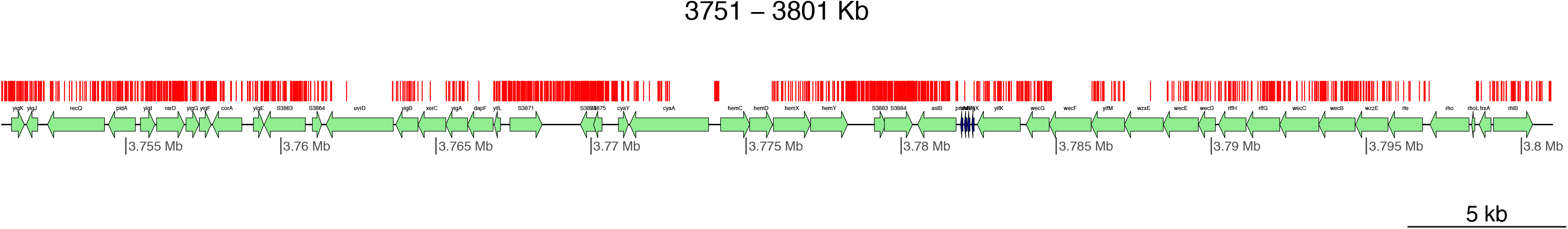
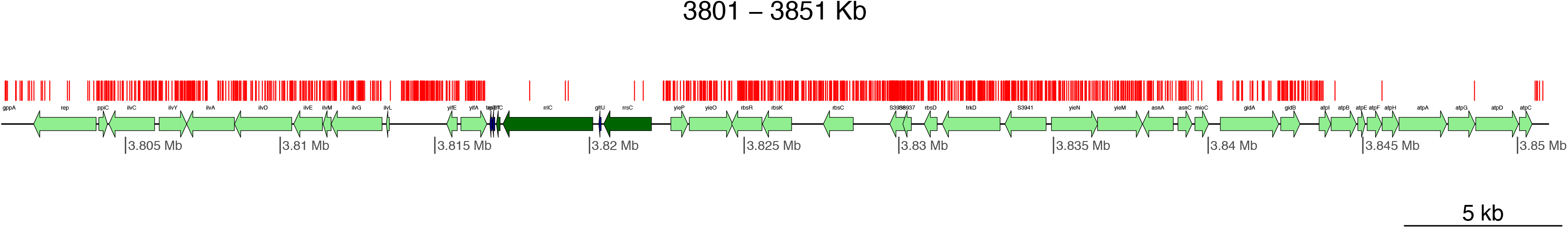
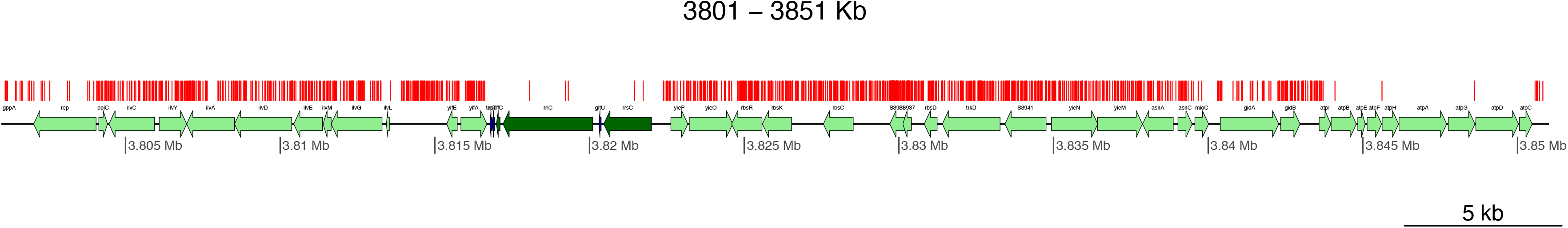
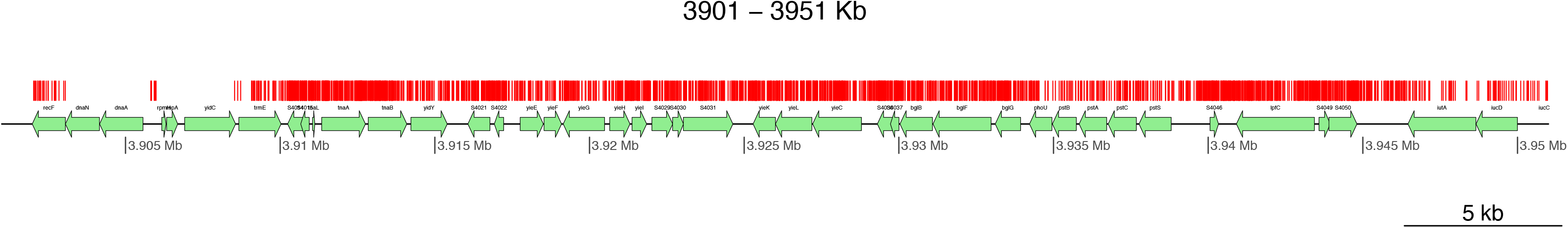
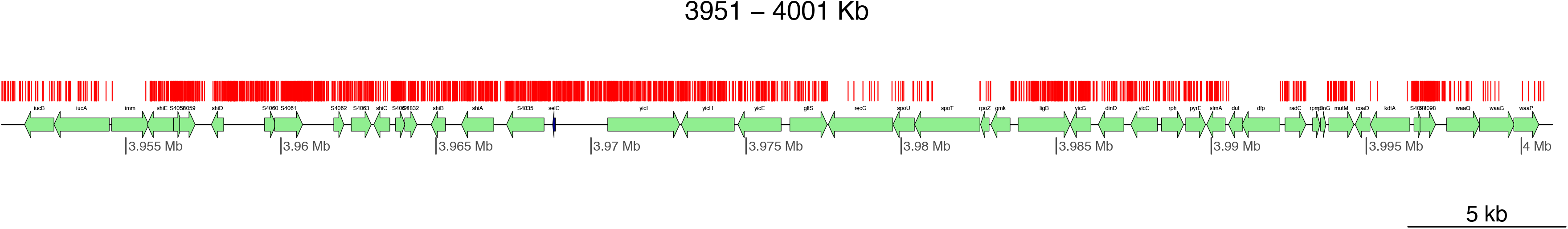
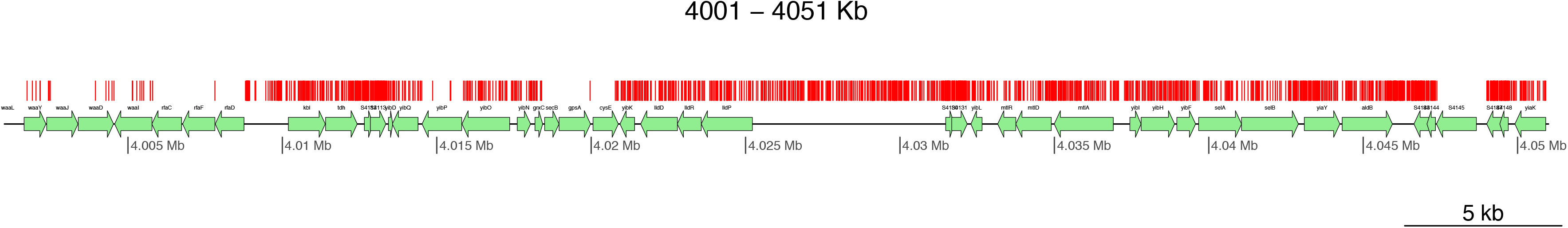
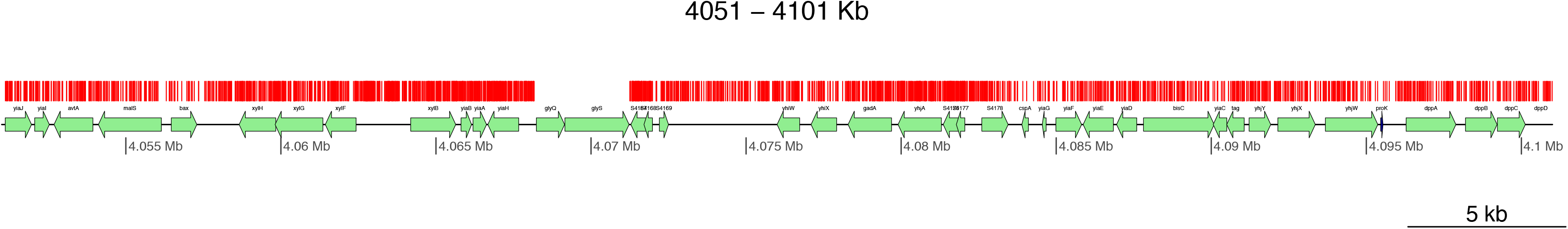
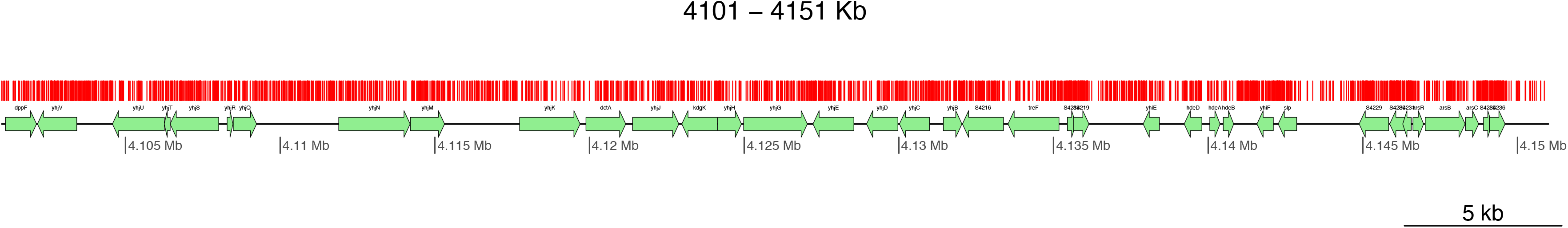
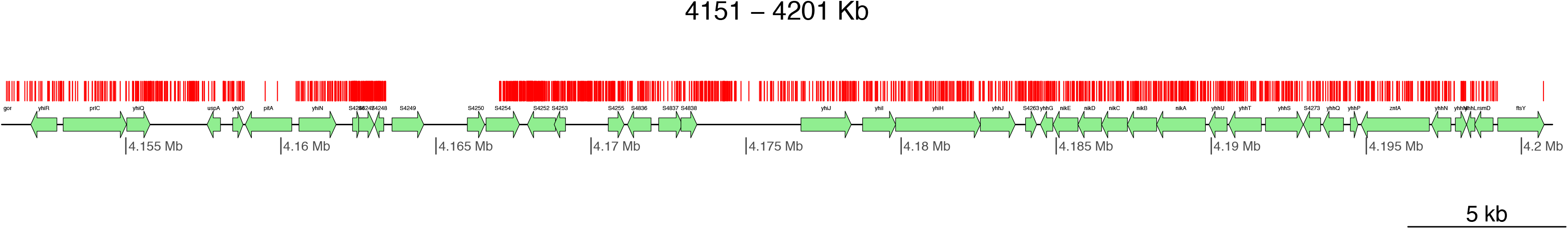
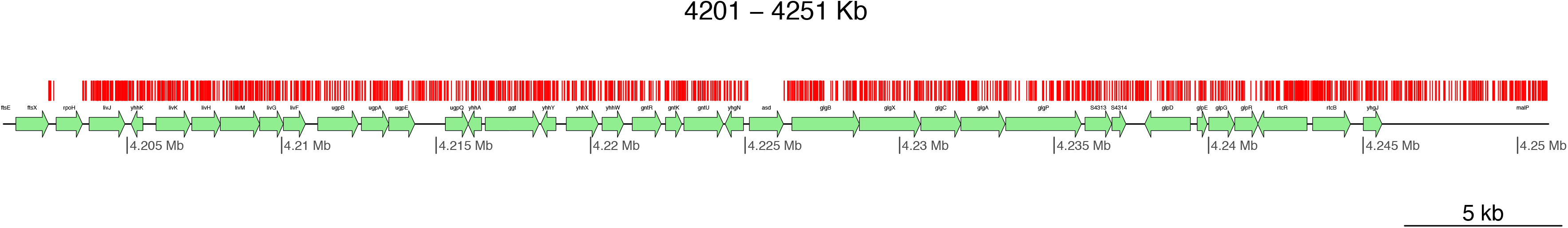
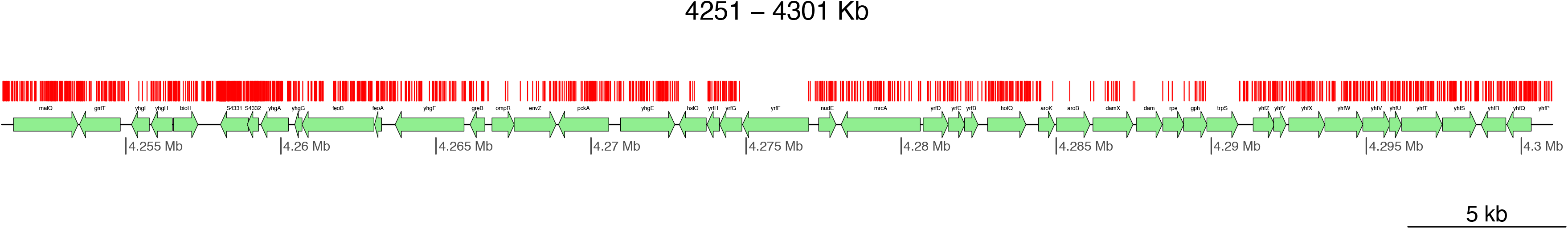
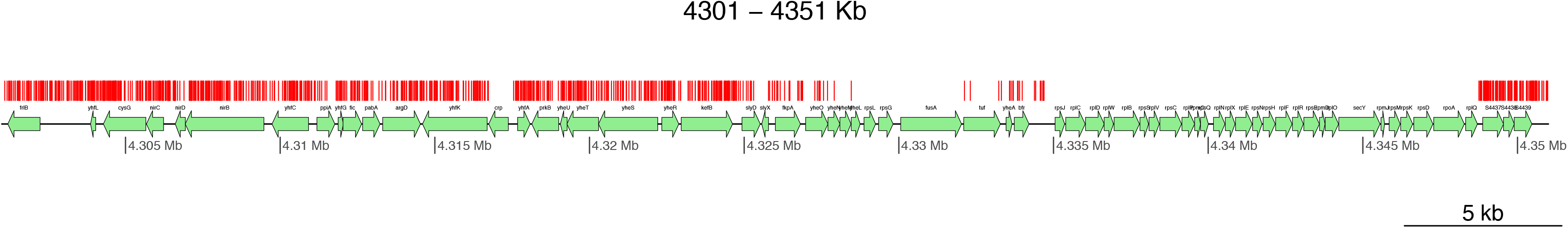
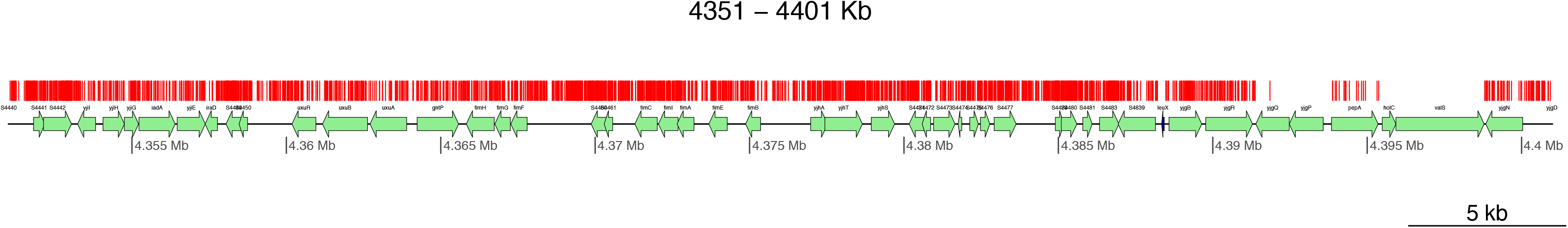
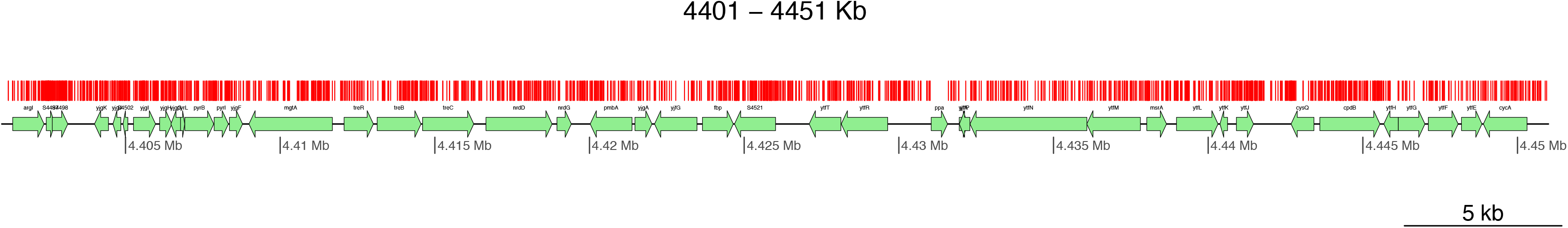
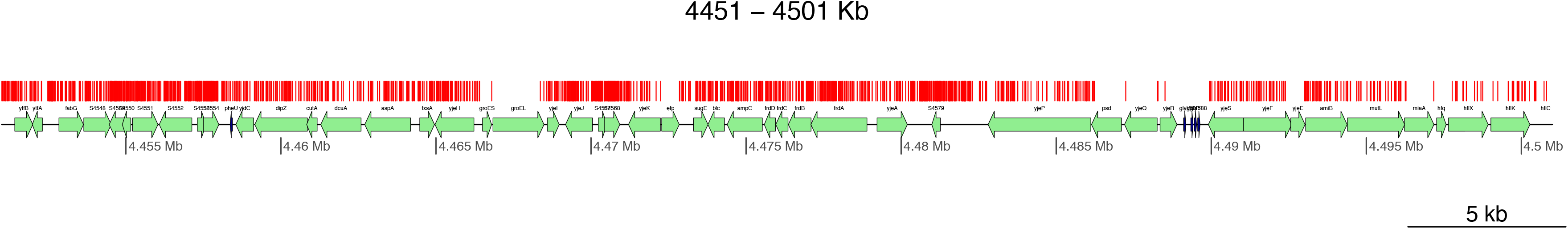
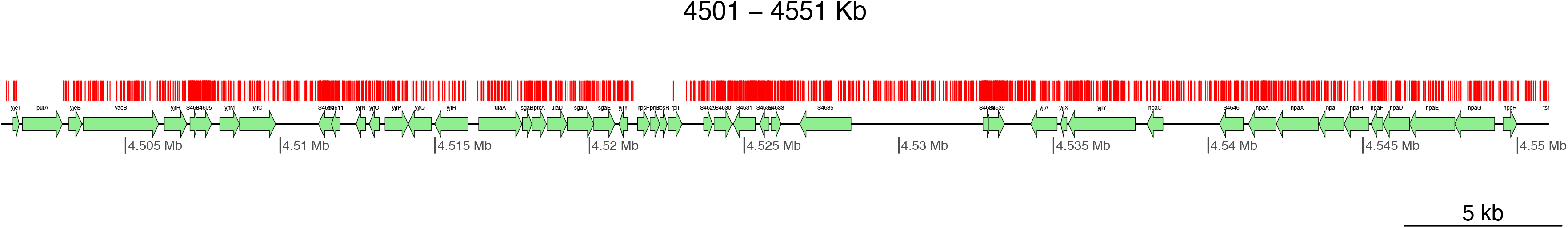
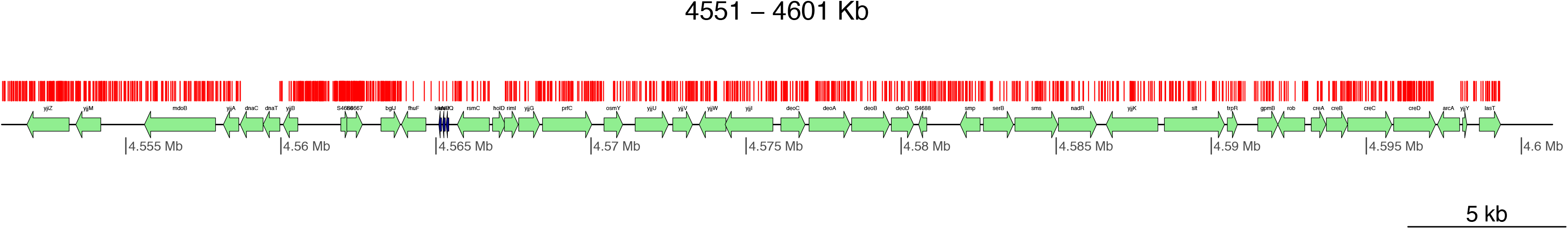
Transposon insertion locations across the entire *Shigella* chromosome. Each insertion site is indicated by a vertical red line. ORFs are indicated in light green; rRNAs in dark green; and tRNAs in dark blue. The annotation is taken from the GenBank sequence of *Shigella flexneri* 2a 2457T.

